# A neural population mechanism for rapid learning

**DOI:** 10.1101/138743

**Authors:** Matthew G. Perich, Juan A. Gallego, Lee E. Miller

**Affiliations:** Department of Biomedical Engineering, Northwestern University, Chicago, IL, USA; Department of Physiology, Northwestern University, Chicago, IL, USA; Department of Physical Medicine and Rehabilitation, Northwestern University, Chicago, IL, USA

## Abstract

Long-term learning of language, mathematics, and motor skills likely requires plastic changes in the cortex, but behavior often requires faster changes, sometimes based even on single errors. Here, we show evidence of one mechanism by which the brain can rapidly develop new motor output, seemingly without altering the functional connectivity between or within cortical areas. We recorded simultaneously from hundreds of neurons in the premotor (PMd) and primary motor (M1) cortices, and computed models relating these neural populations throughout adaptation to reaching movement perturbations. We found a signature of learning in the “null subspace” of PMd with respect to M1. Earlier experiments have shown that null subspace activity allows the motor cortex to alter preparatory activity without directly influencing M1. In our experiments, the null subspace planning activity evolved with the adaptation, yet the “potent” mapping that captures information sent to M1 was preserved. Our results illustrate a population-level mechanism within the motor cortices to adjust the output from one brain area to its downstream structures that could be exploited throughout the brain for rapid, online behavioral adaptation.

## Introduction

A fundamental question in neuroscience is how the coordinated activity of interconnected neurons gives rise to behavior, and how these neurons adapt their output rapidly and flexibly during learning to adapt behavior. There is considerable evidence that learning extended over days to weeks is associated with persistent synaptic changes in the cortex^1–3^. Yet, behavior can also be adapted much more rapidly: motor errors can be corrected on a trial-by-trial basis^4^, and sensory associations can be learned even following a single exposure^5^. Furthermore, in the motor system there appear to be constraints on the types of motor learning that can occur rapidly. In a brain-computer interface experiment, monkeys had difficulty learning to control a computer cursor when a novel control decoder required that they alter the natural covariation among the recorded neurons^6^. There is evidence that such covariance structure relates to synaptic connectivity^7,8^, which may not be readily modified on short time scales (i.e., seconds to minutes)^2,5,9^. Together, these observations suggest that changes in motor cortical structural connectivity may not be the primary mechanism governing changes in neural activity during short-term behavioral adaptation.

To achieve skilled movements, sensory input must be combined with the internal cortical computations needed to achieve the motor action and transformed into a plan executed by the motor cortices^10^. Behavioral adaptation can thus be achieved by adapting this association between sensory input and the motor plan. Dorsal premotor cortex (PMd) is ideally situated to perform the re-association required for short-term learning. PMd is intimately involved in movement planning^11^, has diverse inputs, and shares strong connectivity with the primary motor cortex (M1)^12^. On the other hand, M1 is the main cortical output to the spinal cord with a role primarily in motor execution^13^. Despite the behavioral observations during BCI learning described above and the apparent association between the neural covariance structure and synaptic connectivity, we cannot discount the possibility that fast connectivity changes between PMd and M1 might underlie the adapted behavior. However, an alternative possibility is that short-term motor adaptation is driven by changes in planning-related computations within PMd that are subsequently sent through a stable functional mapping to M1, without any changes in neural covariance and perhaps even synaptic connectivity. At present, such a mechanism has not been described.

Recent work studying population activity during motor planning provides a possible explanation. Using dimensionality reduction methods, the activity of hundreds of motor cortical neurons can be represented in a reduced-dimensional “manifold” that reflects the covariance across the neuronal population^14,15^. When viewed in this way, planning activity in the motor cortices has shown that this neural manifold can be separated into subspaces that are “output-potent” or “output-null” with respect to downstream activity^16^. The potent subspace captures activity that maps functionally onto downstream signals, for example EMG. In contrast, the null subspace captures activity that can be modulated without directly affecting the downstream targets^17^. It appears that PMd planning-related activity remains largely within the null space, seemingly converging toward an “attractor” state that sets the initial conditions necessary to initiate a particular movement^16,18,19^. Proximity to this attractor state is correlated with both reaction time and movement kinematics^20,21^. In our experiment, we applied this framework to study PMd and M1 activity during short-term motor learning. Our results illustrate a novel mechanism within PMd by which the brain rapidly adapts behavior by exploiting the output-null subspace to develop altered planning activity. This mechanism could be thought of as the equivalent of modifying the attractor state to produce the desired action^22^. Ultimately, the adapted initial conditions for that action would be carried to M1 for execution via a fixed output-potent mapping.

### Possible mechanisms underlying adaptive neural activity changes in PMd and M1

In our experiment, we recorded simultaneously from electrode arrays implanted in both M1 and PMd as macaque monkeys learned to make accurate reaching movements that were perturbed either by “curl field” (CF) forces applied to the hand^23–25^, or by a rotation of the visual feedback (VR)^26,27^. We developed an analytical approach based on functional connectivity to understand adaptation-related changes in the activity of M1 and PMd and distinguish the role of the null and potent subspaces within PMd throughout this adaptation process. We trained computational cortico-cortical models (Figure 1) to predict the spiking of single M1 and PMd neurons based on the activity of their neighbors, both local (same recording array) and distant (array in different brain area). Using these models, we described statistically the functional connectivity governing the interactions of these cortical areas. We then tested these models throughout motor adaptation to explore three fundamental questions: 1) does the connectivity within M1 and PMd change during adaptation? 2) does the connectivity between the PMd output-potent activity and M1 change? 3) can evolving output-null planning activity in PMd explain behavioral adaptation? Any model that fails to predict throughout adaptation suggests that the particular modeled connectivity has changed. Within this framework, we can make four specific hypotheses that could explain the modulation of M1 firing rates during adaptation (Figure 1). Note that these four possibilities are not mutually exclusive.

#1) The functional connectivity within the local population of neurons can change, preventing the corresponding model from extrapolating throughout learning (Hypothesis #1 in Figure 1). Intuitively, this means the same presynaptic neural spiking causes a different output in the postsynaptic cell.
#2) A change in motor planning within the null subspace allows PMd to change its potent output to M1, which is ultimately sent through an unchanged mapping captured by the potent space. Here, the potent activity should continue to predict the changes in M1 spiking, but the null activity would be unable to (Hypothesis #2).
#3) A change in the mapping between PMd and M1 will not impact the ability to predict local neurons within either PMd or M1, but will prevent both potent and null PMd activity from predicting M1 (Hypothesis #3).
#4) Lastly, we consider the case that adaptation occurs upstream of PMd, and that the observed activity changes in M1 and PMd are solely in response to altered input. In this case, all models should predict well throughout adaptation (Hypothesis #4).

**Figure 1.**
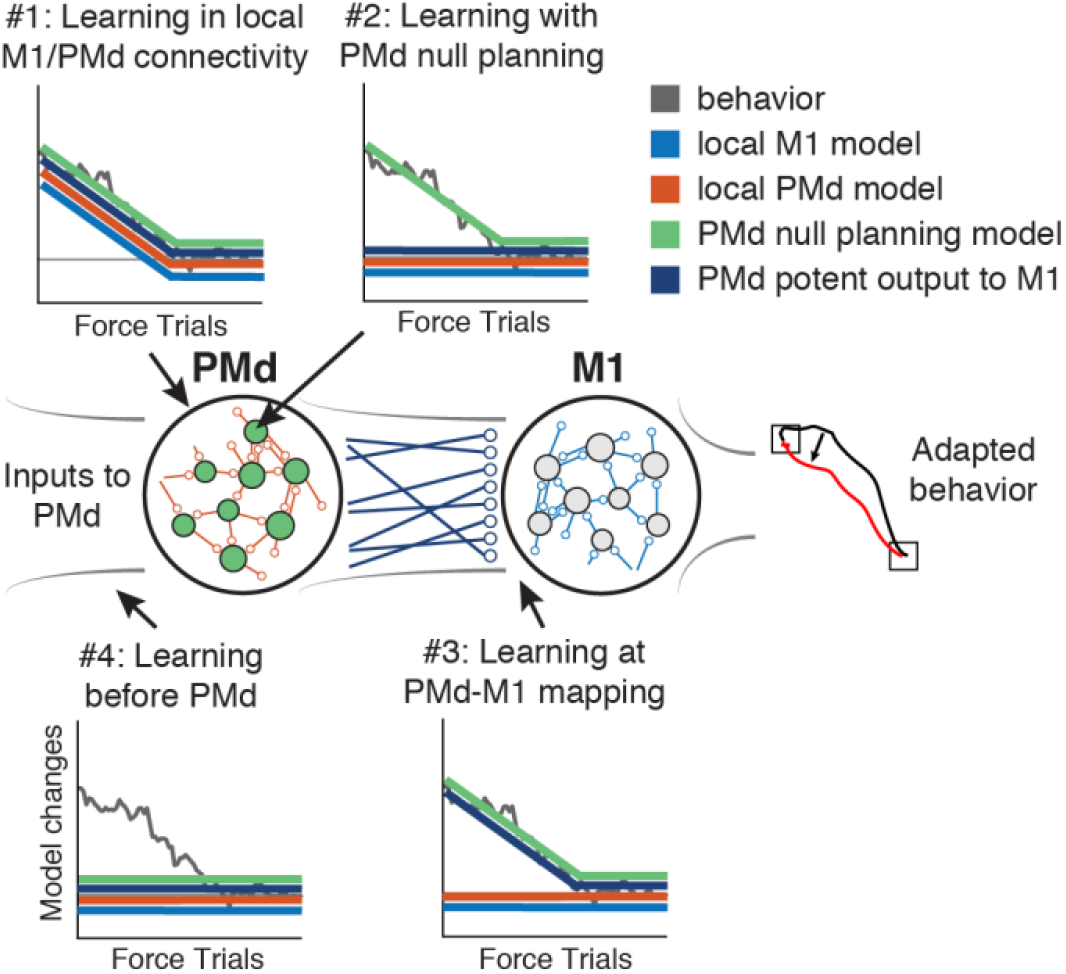
Possible mechanisms of motor cortical activity changes underlying behavioral adaptation. Assuming a simplified hierarchical model where M1 responds to inputs from PMd, we consider four models describing the functional connectivity within M1 (blue), within PMd (orange), between the output-potent activity of PMd and the M1 neurons (dark blue), or the output-null activity of PMd and M1 (green). Our experiment studies the generalization of such models during adaptation. Each inset plot shows the time-course of changes in the behavior (gray line) and the predicted change in modeled connectivity (colored lines). We then identified four hypotheses that could explain the change in firing rates of M1: 1) the local functional connectivity could change, causing all four models to change as behavior adapts; 2) learning could arise from changes in planning-related computations in PMd that are sent to M1. Here, the PMd outputs should predict changes in M1 (dark blue); 3) the mapping between PMd and M1 could change, which would not impact the within-area models (blue and orange) but would prevent the PMd to M1 models from generalizing; 4) learning could occur independently of M1 and PMd, which would not require a change in any of the models describing this circuit.

In the following analyses, we used these cortico-cortical models to explore each of the hypotheses describing short-term adaptation to CF and VR perturbations. We first show that functional connectivity between neurons *within* each area were preserved throughout adaptation to the CF and VR perturbations. We then present evidence that planning activity within the null subspace of PMd plays a direct and unique role in enabling rapid adaptation to the curl field, but not the VR. These results suggest that the CF-related adaptation corresponds to changes in output-null planning activity within PMd, as described by Hypothesis #3 above, while VR adaptation would be mediated by inputs from other brain areas (Hypothesis #4).

## Results

### Curl field adaptation involves widespread, complex changes in firing rate across the motor cortices

We performed experiments to study the functional connectivity relating distinct motor and premotor cortical populations during motor learning, and whether output-potent and output-null activity in premotor cortex enables adapted motor planning. We trained two rhesus macaque monkeys to perform an instructed-delay center-out reaching task (Figure 2a) and implanted 96-channel recording arrays in M1 and PMd (Figure 2b). Each experimental session consisted of three behavioral epochs, beginning with reaches with no perturbation (Baseline) before the monkeys were exposed to the CF in the Force epoch, or the Rotation epoch of the VR experiments. The CF altered movement dynamics and required the monkeys to adapt to a new mapping between muscle activity and movement direction^28,29^ in order to make straight reaches. The VR preserved the natural movement dynamics, but offset the visual cursor feedback by a static 30 deg rotation about the center of the screen. We fi rst consider the CF experiments. Within each session, the monkeys exhibited large errors upon exposure to the CF, which were gradually reduced until behavior stabilized (Figures 2c,d).

**Figure 2.**
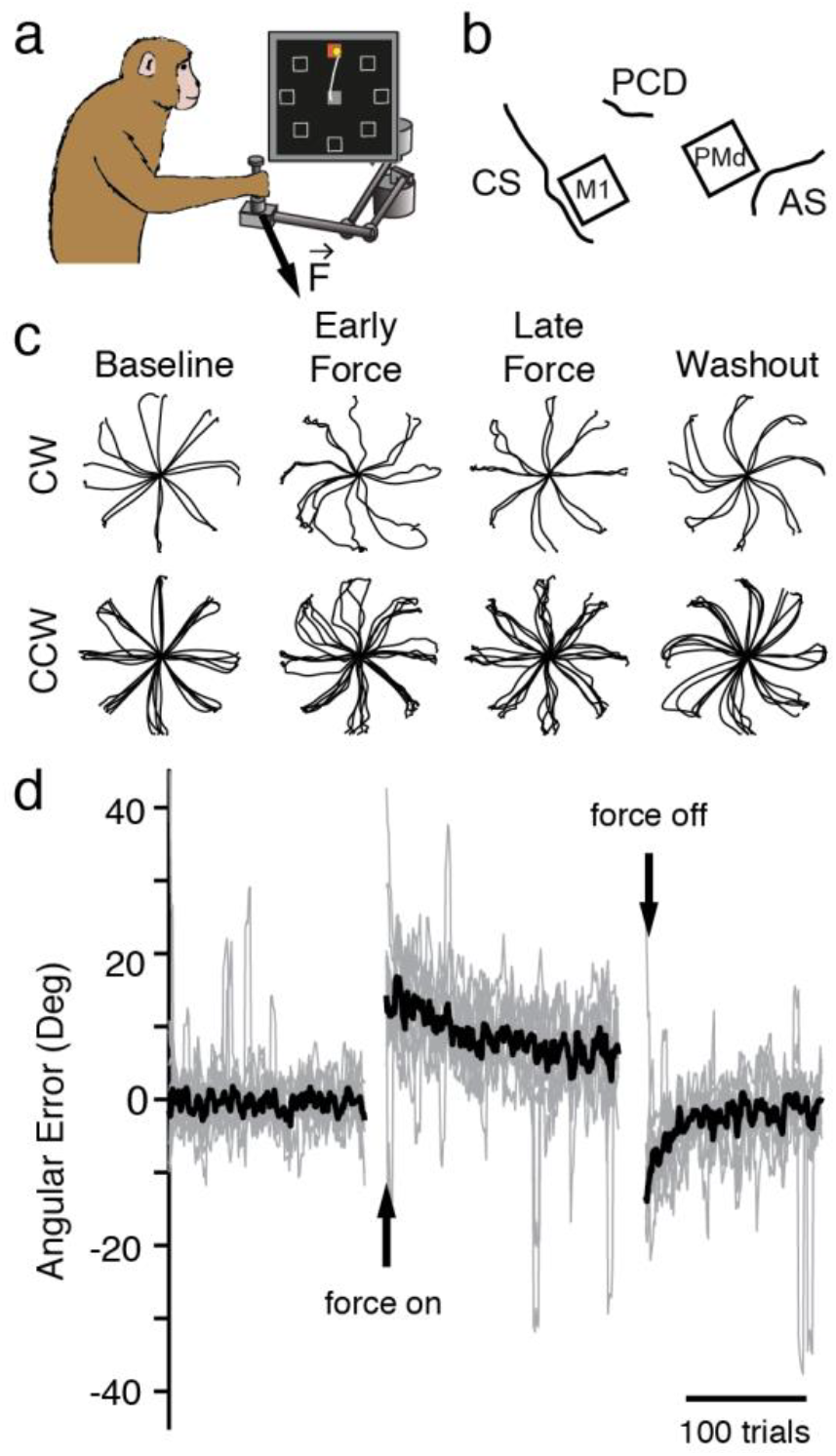
Curl field task. a) Monkeys performed a standard center-out task with a variable instructed delay period following cue presentation. b) We recorded from populations of neurons in M1 and PMd using implanted electrode arrays (right; CS: central sulcus, PCD: pre-central dimple, AS: arcuate sulcus). c) Position traces for the first reach to each target (8cm distance) from two sessions with a clockwise CF (top row) and five sessions with a counter-clockwise CF (bottom row). Sessions from both monkeys are included. d) Error in the takeoff angle for all sessions (light gray lines), with the median across sessions shown in black. Gray traces were smoothed with a 4-trial moving average to reduce noise while preserving the time course.

Further evidence of adaptation was revealed by behavioral after-effects (mirror image errors) that occurred upon return to normal reaching during the Washout epoch (Figure 2d). During Baseline reaching, M1 and PMd neurons exhibited diverse firing rate patterns, as observed in prior studies^30^; a small number of examples are shown in Figures S1 and S2b. At the onset of Force, the firing rate patterns of most neurons changed throughout adaptation along with the movements (Figure S1, S2a,b). Similar changes in firing rate have been previously described in the motor cortex during CF adaptation^24,25,28^. With this study, we explored several competing hypotheses, described above, to explain the underlying cause of these diverse adaptation-related changes in single neuron firing within M1 and PMd.

### Models to study adaptation-related changes in the functional connectivity within or between cortical populations

We designed an analysis to test the above hypotheses using Linear-Nonlinear Poisson Generalized Linear Models (GLMs) to predict the spiking of individual neurons based on the activity of the remaining neurons (Figure 3a; see Methods)^31,32^. Once trained, these models can be used to predict neural spiking during behavioral adaptation even on single trials. We tested whether these models could generalize throughout the Force epoch while behavior changed. We assessed model performance using a relative pseudo-R^2^ (rpR^2^) metric^33^, which quantified the improvement in performance due to the neural covariates above that of the reach kinematic covariates alone (Figure S5a; Methods). Including kinematic covariates helped to account for the linear shared variability related to the executed movement, while leaving the unique contributions of individual neurons (see Discussion and Methods). We separated the adaptation epochs (Force of Rotation) into two sets of trials: training trials taken from the end of the epoch when the behavioral adaptation had stabilized, and testing trials taken from the beginning of adaptation, during rapidly changing behavior. We asked whether our models predicted neural activity throughout adaptation, i.e. whether performance was as accurate for the testing trials as for the training trials (Figure S5d). Failure to generalize indicated that the functional connectivity was altered.

**Figure 3.**
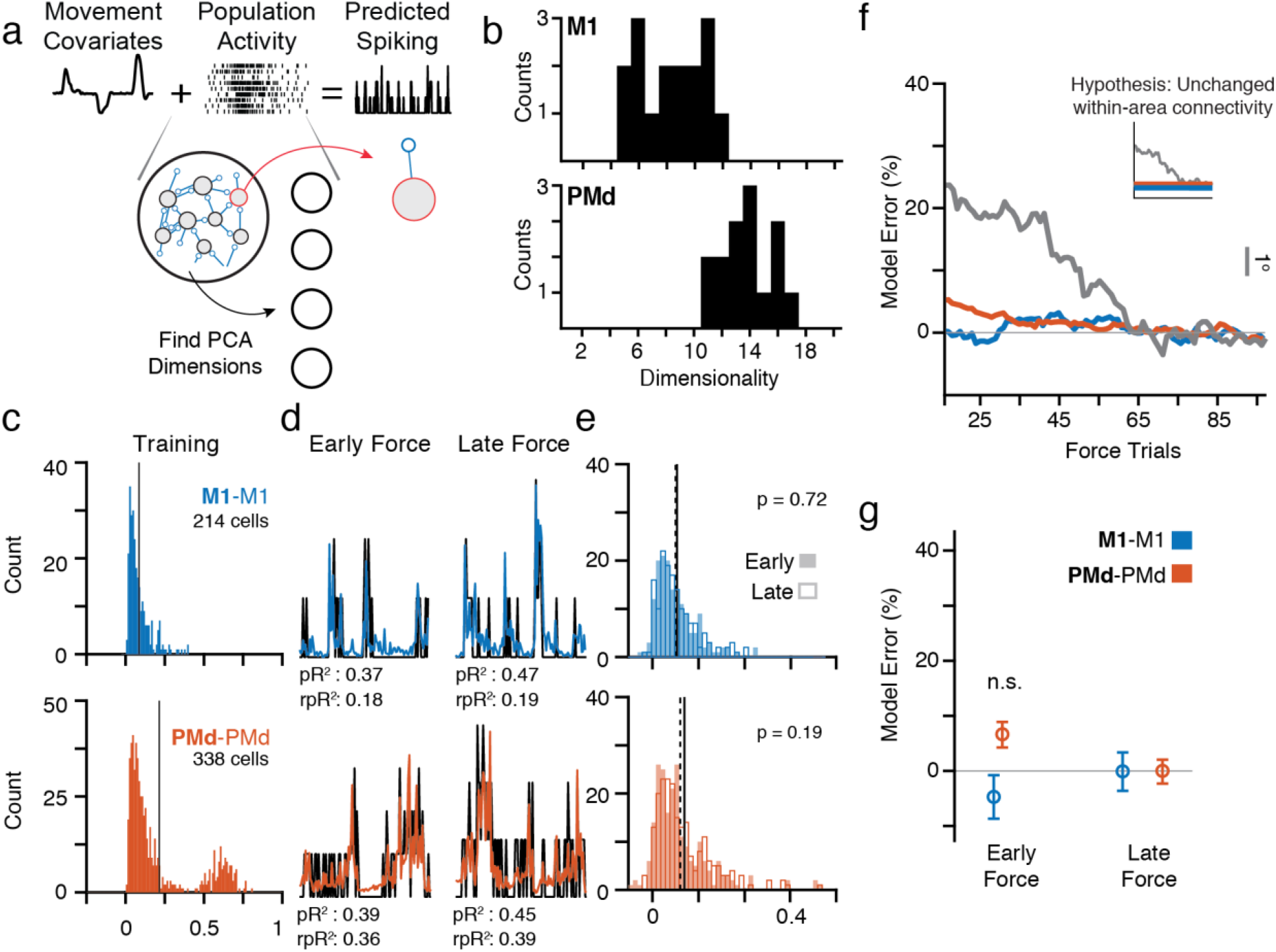
Predicting neural spiking with GLMs. a) We trained GLMs using movement kinematics as well as components of population activity within an area (**M1**-M1, blue, and **PMd**-PMd, orange) to predict the spiking of a left-out single neuron (see Methods). b) Histogram of dimensionality estimates for M1 (top) and PMd (bottom) manifolds across all sessions from both monkeys. PMd activity was consistently higher-dimensional than M1. c) We trained GLMs using data from the end of Force after behavior had stabilized and tested them for generalization throughout the initial phase of adaptation, beginning at the first CF trial (Early) and tested once behavior plateaued (Late). The left histograms show the distribution of rpR^2^ for all cells (pooled across nine sessions) with significant GLM fits (Figure S5; see Methods) based on a cross-validation procedure in the training data from the end of Force. d) Spiking for representative neurons (black) and model predictions (colors) during three Early ((highest behavioral error) and Late (lowest error) adaptation trials. e) Histograms of rpR^2^ values for predictions in the Early and Late blocks. We compared Early (hollow bars; mean: dashed line) and Late (filled bars; mean: solid line) Force trials. **M1**-M1 and **PMd**-PMd had similar distributions during Early and Late. f) Time course of model performance changes. Predictions were made for individual trials, and then smoothed with a 30-trial window (see Methods). Plotted data indicate the mean across all neurons. Trial-to-trial behavioral error processed with the same methods is overlaid in gray and scaled vertically for comparison (gray scale bar on left). The inset replicates our hypothesis prediction from Figure 1. g) Percent error in model performance during the Early and Late adaptation trials as in Panels d and e. No effects were significant (two-sample t-test).

We used the low-dimensional latent activity in PMd and M1 obtained by principal component analysis (PCA)^14,15^ as inputs to the models. This approach implicitly assumes that the neural population covariance matrix (i.e., the neural manifold) is unchanged throughout each experimental session. To confirm this, we compared the covariance between each unit before, during, and after learning (see Methods) and observed no change in covariance throughout the entire experiment (Figure S2; for a single session: Pearson’s correlation with Baseline values r ≥ 0.85 for M1 and r ≥ 0.93 for PMd). Across all sessions, the differences in covariance strongly resembled those obtained from random subsamples of Baseline data for which the covariance structure should be unchanged (Figure S3a). These results show that although neurons in M1 and PMd change their activity to compensate for the curl field, the underlying population covariance remains unchanged.

In order to determine an appropriate number of principal components to use as inputs, we estimated the dimensionality of the manifold that captured the neural population activity (see Methods). In brief, the dimensionality was defined as the number of principal components whose explained variance exceeded a threshold determined by the variance of noise (defined to be variability across trials)^34^. The PMd manifold consistently contained roughly twice the dimensionality as the manifold of M1 (Figure 3b, see Methods), a difference that was robust to the number of recorded neurons (Figure S4d). Thus, for M1 and PMd we used the first eight and sixteen components, respectively, though our results were qualitatively unchanged within a reasonable range of values. Note that the higher dimensionality of PMd necessarily leads to the existence of potent and null subspaces in PMd activity with respect to the M1 latent activity, as hypothesized above.

### Curl field adaptation is not associated with changes in functional connectivity within M1 or PMd

First, we used GLMs to assess whether curl field adaptation was associated with functional connectivity changes within local neural populations (Hypothesis #1 above). To study the functional connectivity within the M1 and PMd populations, we trained two models: **M1**-M1 predicted single M1 neurons from the M1 population activity and **PMd**-PMd predicted PMd neurons from the PMd population. To quantify the model accuracy, we calculated rpR^2^ for each GLM in the “Early” and “Late” blocks of our testing trials taken from the beginning of the Force epoch. We defined the “model error” to be the change in rpR2 normalized by its cross-validated performance (see Methods). We tested whether predictions in the Early block were significantly worse than the Late block, an indication that the models failed to generalize. We set a stringent significance threshold of p = 0.01 for all statistical comparisons. Note that if adaptation affects the accuracy of the models, the highest model error is expected to be in the Early block since it is the furthest removed in time from the training trials. We found that both within-area models (**M1**-M1, **PMd**-PMd) generalized well, predicting the adaptation-related changes in single-cell spiking nearly equally well in both blocks (Figure 3e,f; Figure S6 shows individual monkeys; Figure S7a,b). As a qualitative test of the relation between the behavioral and neural changes during adaptation, we smoothed the rpR^2^ of single-trial GLM predictions and compared their time course to that of the changing behavioral error (Figure 3f; Figure S6 shows individual monkeys). Neither model changed significantly during adaptation (Figure 3g). Based on the good generalization of the GLMs during learning and the unchanging covariance structure (Figures S2, S3), learning is unlikely to arise from changes in the connectivity within M1 or PMd (Figure 1).

### Output-potent activity in PMd consistently predicts M1 neurons across curl field adaptation

We next adapted the models to test whether curl field adaptation corresponded to a change in the output-potent mapping between PMd and M1 (Hypothesis #2 above), or in the output-null computations performed by the PMd population and that seem to capture movement preparation^16,17^ (Hypothesis #3). The analysis of activity in the null and potent subspaces described above provides an elegant framework to separate putative outputs to M1 from the null space putatively comprising the planning computations within PMd (as proposed in Figure 1). We computed the null and potent activity of PMd, defined as the projections of the PMd latent activity into the respective subspaces (see Methods), and used this activity as the inputs for two additional GLMs: **Null**-M1 and **Pot**-M1 (Figure 4b). Importantly, we computed the null and potent subspaces during Baseline, to avoid any possible bias resulting from adaptation-related changes in neural activity during the Force epoch. If the mapping between PMd and M1 changes (Hypothesis #3), **Pot**-M1 will fail to predict M1 spiking during adaptation. Alternatively, if there is a change in the output-null computations reflecting an altered motor plan (Hypothesis #2), then **Pot**-M1, but not **Null**-M1, predicts M1 spiking.

**Figure 4.**
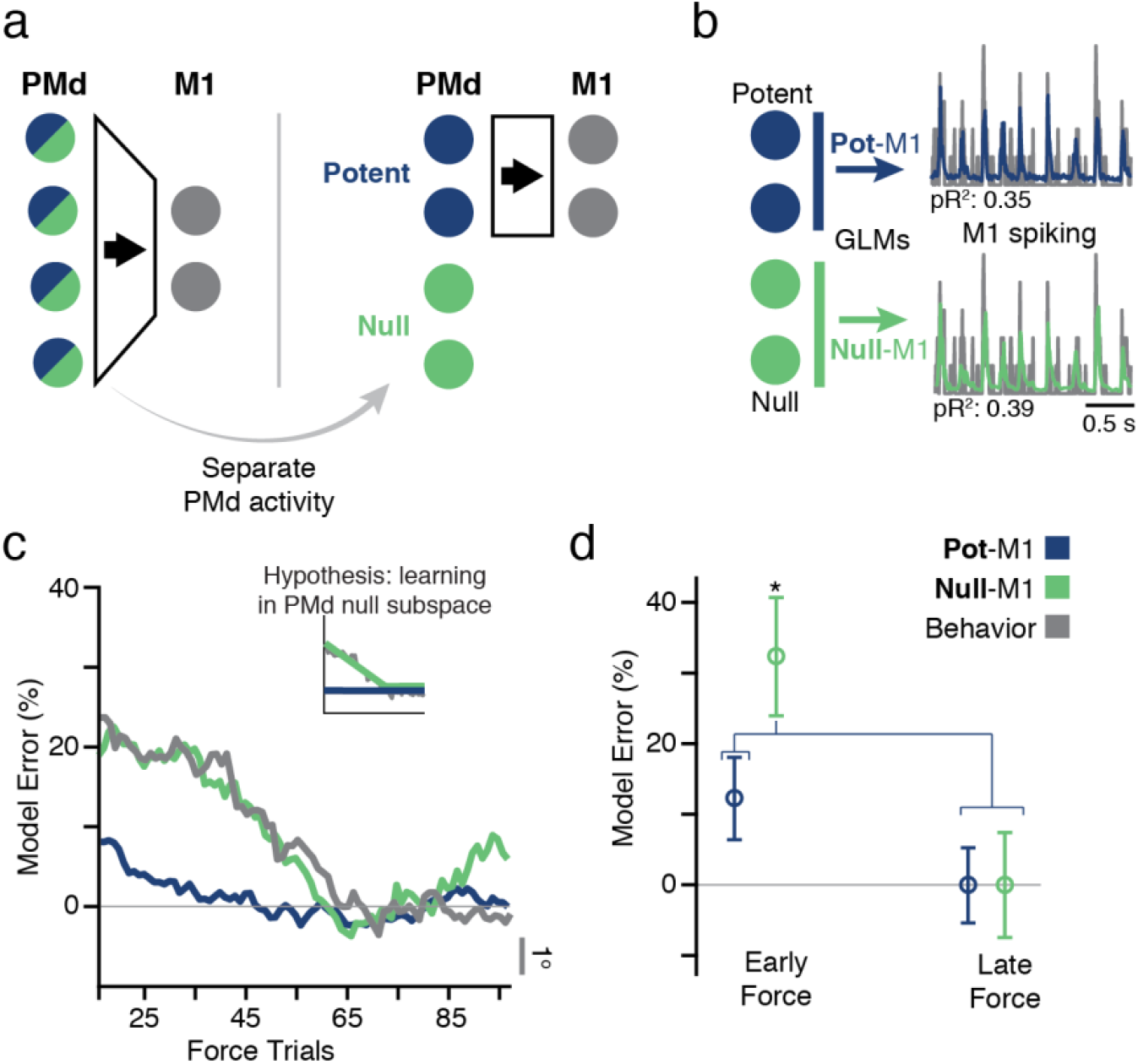
Predicting M1 spiking from PMd potent and null subspaces. a) Hierarchical schematic of motor planning in PMd and M1. The PMd population activity is higher-dimensional, giving extra degrees of freedom at the input to M1 (Figure 3b). We devised an analysis to separate the putative PMd outputs to M1 from the other functions of the population. The former comprises the potent subspace, while the latter reside in the null subspace. b) The potent and null activity of PMd were used as the inputs to a GLM model to predict M1 spiking. c) The time course of model performance for **Pot**-M1 and **Null**-M1 for all sessions. Plots formatted as in Figure 3f. Gray line is the same behavioral error plotted in Figure 3f. Interestingly, the **Null**-M1 error alone paralleled the time course of behavior. d) The bar plot compares model error performance during early and late trials with potent (**Pot**-M1) and null (**Null**-M1) activity. The performance of **Null**-M1 was significantly decreased during adaptation (asterisk indicates significant difference at p < 0.01, two-sample t-test).

Both models had similar accuracy when evaluated within the end-of-Force training trials using ten-fold cross-validation (Figure S5e; see Methods). During learning, however, the time-courses of the prediction accuracy from the potent and null activity were quite different from each other. **Pot**-M1 made accurate predictions in Early Force as well as Late Force (Figure 4d, blue) despite the concurrent behavioral changes. However, predictions of M1 spiking from the PMd null space were significantly worse in the Early block than the Late block (Figure 4d, green). Therefore, curl field motor adaptation is paralleled by changes in the null subspace activity of PMd, while preserving an unchanged output-potent mapping between M1 and PMd (Hypothesis #2 in Fig. 1).

We performed several controls to validate these results. The difference between the models could not be trivially explained by our choice of dimensionality, since we varied the number of principal component inputs across a broad range and observed similar results (Figure S7e). We also developed an additional model, **PMd**-M1, which used all principal components from PMd as inputs. Since **PMd**-M1 contained all of the Potent and Null subspace activity, we expected that, similarly to **Null**-M1, it would fail to generalize during adaptation. Indeed, its accuracy changed progressively during adaptation (Figure S6, pink), with a time-course very much like that of behavior. As a final control, we repeated the GLM analyses using the spiking activity of all single neurons in M1 or PMd as inputs, as opposed to the lower-dimensional latent activity (Figure S7c,d). As expected, the results were very similar since the latent activity captured the dominant patterns of the single neuron spiking.

An important hallmark of motor adaptation is the behavior during Washout: brief mirror-image aftereffects of the errors are seen early in adaptation^23,27^. If the inaccurate **Null**-M1 predictions were a consequence of motor adaptation, we hypothesized that the mapping between null activity in PMd and M1 spiking would return to Baseline during Washout. To test this hypothesis, we trained GLMs using Baseline data and predicted the early Washout trials – conditions which, although separated widely in time, had the same dynamic environment but a very different sta te of adaptation. We found that **Null**-M1 generaliz ed poorly in the earliest trials, then rapidly improved with de-adaptation as null activity began to return to Baseline. There was no such effect for the **Pot**-M1 model, which accurately predicted both early and late Washout trials (Figure 5). The stability of Pot-M1, and the corresponding change in Null-M1, match Hypothesis #2 in Figure 1. Therefore, we conclude that there exists a direct mapping between PMd and M1 that persists unchanged throughout short-term motor adaptation to a curl field, while the evolving null latent activity within PMd changes in a way that drives behavioral adaptation.

**Figure 5.**
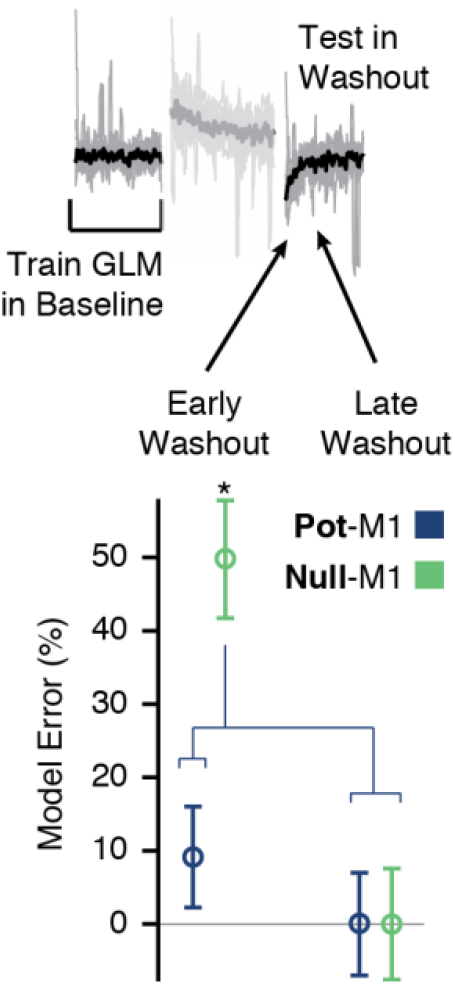
GLM performance during de-adaptation in Washout. We asked whether de-adaptation following the removal of the CF triggered a return to the Baseline **Null**-M1 model. We trained GLMs using Baseline trials and asked how well they generalized in Washout. We found that for **Null**-M1 only, performance was worse in the first eight trials compared to eight trials taken later in Washout. This result supports the idea that the Baseline model was restored in the later trials. Asterisks indicate significant differences at p < 0.01 (two-sample t-test).

### Evolving changes in PMd output-null planning activity lead to adapted motor output

We next investigated the nature of the activity changes in the null and potent PMd subspaces during CF adaptation. To better visualize the changes in latent activity, we plotted activity in the single leading axis of the potent and null subspaces for movements to a single target during a Baseline reach (black) and throughout adaptation (blue shades) (Figure 6a). During behavioral adaptation, potent and null activity both progressively deviated from that of baseline. In this simple example, activity preferentially changes along the output-null axis prior to the go cue, before the activity begins to modulate in the output-potent axes during the subsequent movement. To quantify this effect, we asked whether the activity changes across all targets and sessions were consistent with the idea that they represented adapted preparatory movement plans. Throughout adaptation, we computed the Mahalanobis distance in the low-dimensional neural manifold at the time of go cue between the neural activity in Force and the corresponding activity in Baseline (Figure 6b). The distance increased monotonically throughout adaptation. Critically, since the increased distance was observed already at the time of the go cue, well before the onset of movement, these changes are suggestive of an adapting motor plan, as we predicted in Hypothesis #2.

**Figure 6.**
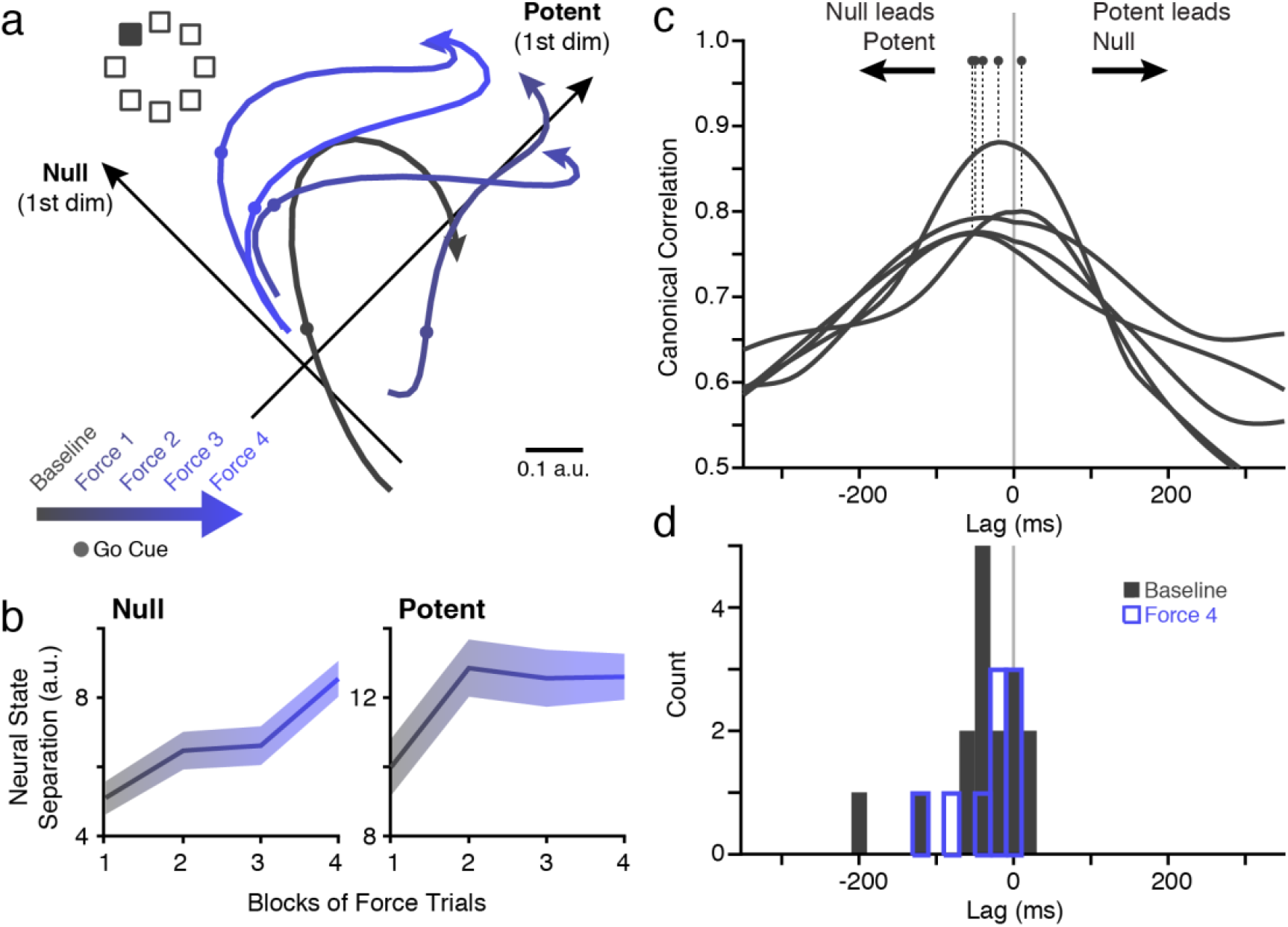
Potent and Null activity during CF adaptation. a) The evolution of potent and null activity during CF adaptation for a single reach direction (filled square) averaged in four blocks of trials. Activity along the first null and potent PMd axes changed with adaptation. Notably, the change between early and late Force were evident at the time of go cue, before movement began (circles). b) Analysis of neural state shows a progressive change in potent and null activity during adaptation. At the time of go cue, we computed the Mahalanobis distance for all target directions, which represents the separation of the instantaneous neural state from that of the Baseline. Plots show mean and s.e.m. across all CF sessions. c) We used Canonical Correlation Analysis (CCA; see Methods) to compare the null and potent activity at different time shifts to identify their relative lag. The first canonical correlation value is shown for five example sessions. Dots indicate the lag with peak correlation. A peak left of zero indicates that null leads potent activity. d) Peak lag across Baseline for all 16 CF and VR sessions (black histogram). Null activity preceded potent by 35 ms on average, and the distribution was significantly negative (t-test; p = 0.02). The distribution obtained from the end of the Force period for the CF sessions (9 sessions) is overlaid in blue. The shift in time did not change during learning.

In the preceding results, we interpret the null activity as the computations necessary to formulate a new motor plan, a plan that can ultimately influence activity in M1. One prediction from this interpretation is that the null activity should precede the potent activity to which it should be linked^16,17^. We estimated the lag between the null and potent activity by computing the canonical correlations at varying time shifts (see Methods). We identified the shift at which the correlation was maximum and found that null activity consistently did precede that of potent (and thus also M1; Figure 6c,d), during both Baseline and at the end of Force, when behavior was fully adapted. This observation, coupled with the GLM results, provides evidence that the changes in null activity could be causally related to the formation and subsequent execution of an adapted motor plan.

### PMd to M1 mapping is unchanged throughout adaptation to a visuomotor rotation

In an important parallel experiment, we asked whether changes between PMd and M1 are a necessary consequence of adapted behavior, or indicative of a specialized role for PMd in the CF task. The same monkeys also learned to reach in the presence of a 30° visuomotor rotation (VR), a static rotation of the reach-related visual feedback (Figure 7a). Considerable evidence suggests that the brain areas involved in adapting to the VR differ from those required for CF^35–37^, and include the parietal cortex, upstream of PMd. If the change in **Null**-M1 mapping during CF adaptation represents updated motor planning within PMd, we hypothesized that VR adaptation might not result in a similar change. Instead, VR adaptation is mediated by processes occurring before the inputs to M1 and PMd (Hypothesis #4, Figure 1).

**Figure 7.**
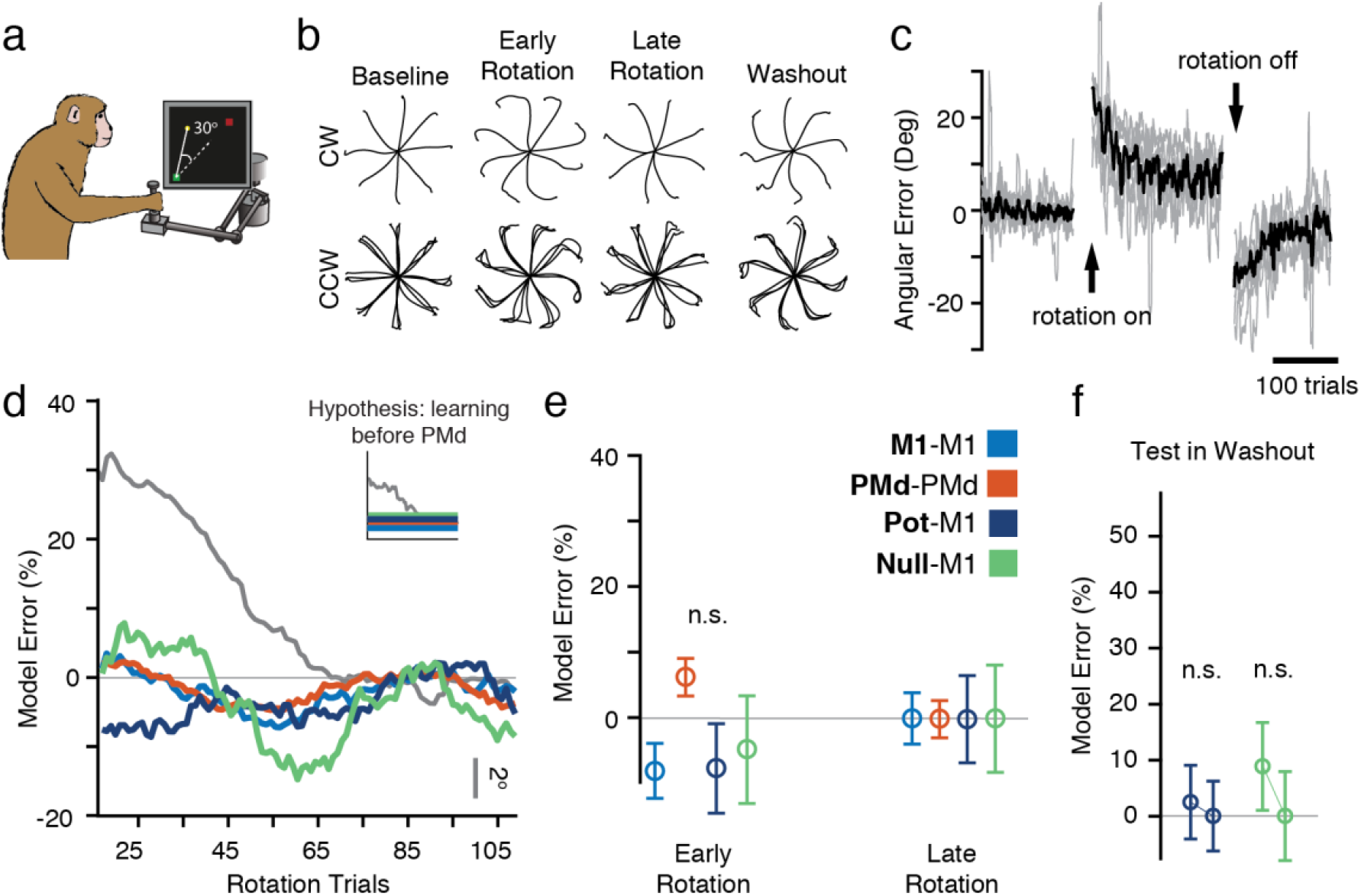
Adaptation during a visuomotor rotation task. a) The same monkeys also adapted to a 30° visuomotor rotation (VR). b) c) Behavioral adaptation traces, plotted as in Panel b of Figure 2, for the VR sessions (one clockwise and four counter-clockwise). c) Same as Panel c of Figure 2 but for the VR sessions, showing behavioral errors similar to those of the CF condition. d) The time course of model prediction error during rotation adaptation. Inset: we hypothesized that compensation for the altered feedback of the VR involved brain structures upstream of PMd. In this case, all three of the GLM models should generalize. As predicted, all GLM models, including Null-M1 generalized well to both Early and Late Rotation trials, despite clear behavioral adaptation (gray). e) Same format as Figure 3g. There were no significant changes throughout adaptation (all p > 0.01, two-sample t-test). f) Washout model performance for the VR task. Methods and data are as presented in Figure 5. We observed no change in performance during de-adaptation.

In many respects, adaptation during the CF and VR sessions was quite similar. Behavioral errors were similar in magnitude and time course (Figure 7b,c), with large initial errors in the Rotation epoch, and strong after-effects in Washout. Single neurons also changed their firing in complex ways (Figure S1b), and the altered activity was observed during pre-movement planning. Additionally, changes in neural activity preserved the population covariance structure (Figure S2b). However, all GLM models, including **Null**-M1, accurately generalized throughout Rotation (Figure 7d,e). Furthermore, we did not see the effect of de-adaptation in the Washout predictions, despite the presence of behavioral after-effects (Figure 7f). Thus, there were no changes in the relationships between the PMd null activity and M1, despite behavioral adaptation and diverse changes in single-neuron activity. This result highlights a fundamental difference in the neural adaptation to these two perturbations, and supports the view that VR adaptation occurs upstream of PMd^35–37^. It also strengthens our conclusions about the CF task: the poor generalization of the **Null**-M1 GLM is not a necessary consequence of changing behavior, but rather captures a previously undescribed mechanism by which the premotor cortex can drive rapid sensorimotor adaptation by exploiting the output-null subspace.

## Discussion

We have reported an experiment designed to help understand how the motor cortex can rapidly adapt behavior. We analyzed the functional connectivity between local and distant neural populations in M1 and PMd during two common motor adaptation tasks and studied PMd activity in output-potent and output-null subspaces. Our analyses revealed several intriguing results: 1) during both short-term adaptation tasks, the functional connectivity within M1 and PMd assessed using GLMs, as well as the neural covariance, was unchanged; 2) the estimated dimensionality of PMd activity was larger than that of M1, suggesting the existence of null and potent subspaces; 3) PMd potent activity had a consistent relationship with M1 throughout adaptation in both tasks; 4) the relationship between PMd null activity and neurons in M1 changed with behavioral adaptation and subsequent de-adaptation in Washout for the curl field task (Figures 4,5,S6), but not visuomotor rotation (Figure 7); 5) pre-movement activity within PMd appears to reflect motor planning within the null space that evolves gradually during adaptation (Figure 6). This study was enabled by recording large neural populations simultaneously from two distinct regions of the motor cortex; none of these observations would have been possible with sequential single-neuron recordings. Our results are inconsistent with the idea that changes in connectivity is a necessary mechanism for short-term motor learning. Instead, our results show that within the constrained, low-dimensional manifold, the cortex is able to utilize the output-null subspace to change its output on a trial-by-trial basis. The use of output-null population activity while maintaining an unchanged output-potent projection to downstream structures, could provide a powerful mechanism underlying a variety of rapid learning processes throughout the brain.

### Interpreting output-null activity in PMd

The relation between the null and potent subspaces in movement execution and motor learning is an intriguing area of inquiry. Two recent studies using natural reaching movements and a BCI paradigm suggest that the appropriate neural state must be established within the cortical null subspace to initiate potent activity and subsequent motor output^16,38^. In our CF experiment, we observed progressive adaptation-related changes in the null activity during the planning period preceding movement (Figure 6), and corresponding changes during the more rapid de-adaptation process in Washout. Critically, the null activity led the potent activity following target presentation. Together with evidence from prior studies implicating null activity in motor preparation^16,17,39^, our results strongly suggest that the evolving activity within the null subspace of PMd serves to set the preparatory state within PMd prior to movement initiation. One possible interpretation is that during movement preparation, an attractor-like state within PMd is altered, in order to establish new initial conditions for the execution of an adapted movement by M1^22^. At the transition between movement planning and initiation, the null and potent subspaces are thought to be transiently coupled, such that the state in the null subspace initiates the cortical dynamics necessary to cause the desired motor output^16,17,40^. The nature of this transition, including the underlying mechanism and its timing, are the subject of ongoing experimental and modeling efforts^40^. Kaufman et al. propose that the transition could be driven by influence from a third area, perhaps a higher cortical area or the basal ganglia^40^. In our experiments, the altered state in the null subspace is used to initiate a new, adapted movement. Importantly, this process could be accomplished without changing either the functional connectivity within either area, or the potent mapping from PMd to M1.

The manifold comprising the activity within PMd was divided into null and potent subspaces relative to the latent activity of M1^16^. We computed the subspaces using data from the Baseline epoch, i.e., independently of the CF trials used for the GLM analysis. By definition, the null activity orthogonal to that of the potent activity and M1 latent activity. An obvious question then, is how it is possible to use GLMs to predict the activity of M1 neurons from null activity in PMd? Although the potent and null subspace activity reside in orthogonal axes, the activity along any two of these axes can be linearly related during behavior since they are constructed from different weighted combinations of the common set of PMd latent activity. This explains why both null and potent activity can be used to predict M1 spiking in the GLMs. Yet, during learning the precise relationship between null activity and M1 appears to evolve with adaptation, while potent activity maintains the same relation with M1 spiking throughout the entire adaptation period. What is the correct way to interpret these activity changes? At any given time, the neural activity can occupy any portion of the potent or null subspaces. Adaptation could result in a change in either the orientation of these subspaces within the higher-dimensional PMd manifold, the trajectory of PMd neural activity within the fixed subspaces, or both (Figure S3c illustrates these distinctions). The similarity of the neural covariance structure (Figure S2, S3) is evidence that the orientation of the M1 and PMd manifolds did not change during adaptation. Instead, activity within these fixed subspaces changed as the monkeys learned.

### Testing the limits of the relations inferred from GLMs

While it is tempting to infer detailed cortical circuitry from the statistical structure in the neural population activity, it is dangerous to do so. The simple neural covariance approach (Figures S2,S3) is particularly far from cortical circuitry, as it is driven to a great extent by the common input received by all the recorded neurons^41^. Furthermore, because many trials of data are necessary to estimate the covariance structure accurately, such covariance approaches are ill-suited to study trial-by-trial adaptive changes. Our GLMs are instead able to predict spiking activity on single trials. Furthermore, the GLMs can make a better estimate of the direct statistical dependencies, which we interpret as functional connectivity, between groups of cells by discounting some of the common input that drives their activity. To achieve this, other studies have incorporated a wide variety of covariates, such as spiking history, the activity of neighboring neurons, expected reward, or kinematics^31,42,43^. We included additional kinematic covariates as inputs to the models to account for the shared inputs to the neurons (see Methods)^31,42–44^, though our results were not qualitatively dependent on the precise set of signals used. Note that our use of kinematic signals in the model does not assume a direct link between neural activity and movement kinematics. Rather, it is an attempt to discount influences in common input across a large number of neurons, which would otherwise obscure subtler, unique influences. Here, this approach revealed a differential effect of adaptation on the functional connectivity between M1 and the null and potent subspaces of PMd.

### Visuomotor rotation and curl field likely employ distinct adaptive processes

The visuomotor rotation experiments provide an intriguing counterpoint to curl field results. Adaptation to the VR is believed to involve neural processes that are independent from those of the curl field^35,45^. Previous evidence from modeling studies has led to the proposition that VR adaptation involves changes in functional connectivity between parietal cortex and motor and premotor cortices^37^. Consistent with that proposal, despite the dramatic changes in PMd and M1 activity during VR adaptation (Figures S1,S2b), all GLM models generalized well: there were no changes in PMd to M1 functional connectivity throughout adaptation (Fig. 6), and we did not observe the effect of de-adaptation in the Washout **Null**-M1 GLM mapping (Figure 7f). We interpret this result to mean that VR adaptation occurs upstream of the inputs to PMd, although the current experiment cannot show this directly. This experiment also serves as an important control for our main result in the CF task, because a motor adaptation process with very similar magnitude of error and time course of adaptation yielded a very different result. An interesting future experiment would be to repeat the GLM modeling analysis using simultaneous recordings from parietal cortex and PMd during VR adaptation. In this case, we would expect to see evolving null subspace activity in parietal cortex similar to that observed in PMd during the CF task.

### Relation between short- and long-term learning

Long-term learning is known to alter connectivity in the motor cortex, resulting in increased horizontal connections^46^ and synaptogenesis^47^. Many have proposed that the brain also uses similar plastic mechanisms to adapt behavior on shorter timescales^24,48^. However, any such changes of behaviorally-relevant magnitude would very likely have impaired the predictions of the **Pot**-M1, **M1**-M1, or **PMd**-PMd GLM models^49,50^. Hence our results suggest that functional connectivity changes within PMd or M1 play at most a minimal role on the time scale of a single experimental session. Our lab has previously found that the relation between evolving M1 activity and the dynamics of the motor output remains unchanged during CF adaptation^28^, with no evidence for adaptive changes in spatial tuning having a time course like that of behavioral adaptation^25^. Therefore, we hypothesized that CF adaptation must be mediated by changes in recruitment of M1 by upstream areas, including PMd. Our new results illustrate how this can occur: in the short-term, PMd could exploit its null subspace to formulate new motor plans reflecting the modified task demands of the CF, which are sent to M1 through a fixed potent mapping.

However, if short-term CF adaptation occurs without connectivity changes in the cerebral cortex, how can we account for the performance improvements that are maintained across sessions^51,52^? The cerebellum has long been implicated in a variety of supervised, error-driven motor-learning problems, including both the CF and VR paradigms explored in this study^36,53–55^. Importantly, the cerebellum is thought to be a site at which inverse internal models of limb dynamics may be learned^56,57^. Furthermore, plastic adaptive processes occur at multiple sites in the cerebellum and over several time scales^58,59^. Perhaps its extensive interconnections with PMd^60^ allow new motor plans developed during CF adaptation to drive an evolving cerebellar internal model^4,36,57^. Other evidence suggests that while these internal models may depend on the cerebellum for their modification, they may actually be located elsewhere^61^. In this case, such connectivity changes might eventually emerge within PMd and even M1 to support the long-term refinement and rapid recall of skills^1,62^.

Wherever their ultimate location, we envision a mechanism whereby the tentative plans progressively developed in something resembling a “neural scratch pad” of cortical null subspaces, ultimately become consolidated in neural structural changes. Analogous population dynamics have been found in prefrontal cortex for decision-making^63^, working memory^34^, and rule-learning^64^, in the motor cortex for movement planning^16^, and in the parietal cortex for navigation^65^ among others^15^. These widespread observations suggest a general mechanism by which the brain may leverage output-null subspaces of cortical activity for rapid learning without the need to engage potentially slow processes to alter the cortical circuitry.

## Acknowledgements

The authors would like to thank Dr. Sara Solla for her helpful discussions in preparing these analyses. J.A.G. received funding from the European Commission (FP7-PEOPLE-2013-IOF-627384). M.G.P. received funding from NIH NINDS T32 HD07418 and NIH NINDS F31 NS092356. This project was additionally funded by NIH NINDS NS053603 and NIH NINDS NS074044.

## Ethical Statement

All procedures involving animals in this study were performed in accordance with the ethical standards of the Northwestern University Institutional Animal Care and Use Committee and are consistent with Federal guidelines.

## Author Contributions

M.G.P. and L.E.M. conceived and designed experiments. M.G.P. conducted experiments and analyzed data. M.G.P, J.A.G., and L.E.M. prepared figures, designed analyses, interpreted results, and wrote the manuscript.

## Methods

All animal-related methods were approved by the Institutional Animal Care and Use Committee of Northwestern University and were consistent with federal guidelines.

### Behavioral task

Two monkeys (male, *mucaca mulatta;* Monkey C: 11.7 kg, Monkey M: 10.5 kg) were seated in a primate chair and made reaching movements with a custom 2-D planar manipulandum to control a cursor displayed on a computer screen. We recorded the position of the handle at a sampling frequency of 1kHz using encoders. The monkeys performed a standard center-out reaching task with eight outer targets evenly distributed around a circle at a radius of 8 cm. All targets were 2 cm squares. The first three sessions with Monkey C used a radius of 6 cm. However, we observed no qualitative different in the behavioral or neural results for the shorter reach distance, and all sessions were thus treated equally. Each trial began when the monkey moved to the center target. After a variable hold period (0.5 – 1.5 s), one of the eight outer targets appeared. The monkey had a variable instructed delay period (0.5 – 1.5 s) which allowed us to study neural activity during explicit movement planning and preparation, in addition to movement execution. The monkeys then received an auditory go cue, and the center target disappeared. The monkeys had one second to reach the target, and were required to hold there for 0.5 s.

In the curl field (CF) task, two motors applied torques to the elbow and shoulder joints of the manipulandum in order to achieve the desired endpoint force. The magnitude and direction of the force depended on the velocity of hand movement according to Equation 1, where 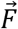 is the endpoint force, *ṗ* is the derivative of the hand position 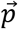, *θ_c_* is the angle of curl field application (85°), and *k* is a constant, set to 0.15 N·s/cm:

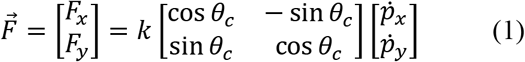

In the visuomotor rotation (VR) task, hand position *p* was rotated by *θ_r_* (here, chosen to be 30°) to provide altered visual cursor feedback 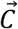 on the screen. The rotation was position-dependent so that the cursor would return to the center target with the return reach:

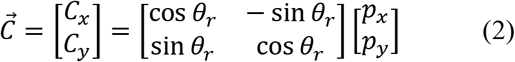

Both the CF and VR perturbations were imposed continuously throughout the block of adaptation trials, including the return to center and outer target hold periods.

Each session was of variable length since we allowed the monkeys to reach as long as possible to ensure that behavior had sufficient time to stabilize, and to allow for large testing and training sets for the GLMs. For the CF sessions, the monkeys performed a set of unperturbed trials in the Baseline epoch (range across sessions: 170 – 225 rewards) followed by a Force epoch with the CF perturbations (201 – 337 rewards). The session concluded with a Washout epoch, in which the perturbation was removed and the monkeys readapted to making normal reaches (153 – 404 rewards). The curl field was applied in both clockwise (CW) and counter-clockwise (CCW) directions in separate sessions, though we saw no qualitative difference between the sessions. Monkey C had three CW sessions and two CCW sessions, while Monkey M had four CCW sessions. For the VR sessions, the monkeys performed 154 – 217 successful trials in Baseline, 219 – 316 during Rotation (either CW or CCW), and then 162 – 348 in Washout. Monkey C performed two CW VR sessions and two CCW sessions, while monkey M performed three CCW sessions. There is considerable evidence that learning can be consolidated, resulting in savings across sessions^51^. In this study, we minimized the effect of savings to focus on single-session adaptation in the following ways. The monkeys: 1) received different perturbations day-to-day, as we alternated between CF and VR sessions, 2) received opposing directions of the perturbation on subsequent days, and 3) often had multiple days between successive perturbation exposures.

### Behavioral adaptation analysis

For a quantitative summary of behavioral adaptation, we used the errors in the angle of the initial hand trajectory. We measured the angular deviation of the hand from the true target direction measured 150 ms after movement onset. To account for the slightly curved reaches made by the monkeys, we found the difference on each trial from the average deviation for that target in Baseline trials. Sessions with the CW and CCW perturbations were similar except for the sign of the effects. Thus, for the behavioral adaptation time course data such as that presented in Figure 2d, we pooled all perturbation directions together and flipped the sign of the CW errors.

### Neural recordings

After extensive training in the unperturbed center-out reaching task, we surgically implanted chronic multi-electrode arrays (Blackrock Microsystems, Salt Lake City, UT) in M1 and PMd. From each array, we recorded 96 channels of neural activity using a Blackrock Cerebus system (Blackrock Microsystems, Salt Lake City, UT). The snippet data was manually processed offline using spike sorting software to identify single neurons (Offline Sorter v3, Plexon, Inc, Dallas, TX). We sorted data from all three task epochs (Baseline, Force or Rotation, and Washout) simultaneously to ensure we reliably identified the same neurons throughout each session. With such array recordings, there is a small possibility that duplicate neurons can appear on different channels as a result of electrode shunting, which would influence our GLM models by providing perfectly correlated inputs for these cells. While such duplicate channels are often easily identifiable during recording, we took two precautionary steps to ensure our data included only independent channels. First, we used the electrode crosstalk utility in the Blackrock Cerebus system to identify and disable any potential candidates with high crosstalk. Second, offline we computed the percent of coincident spikes between any two channels, and compared this percentage against an empirical probability distribution from all sessions of data. We excluded any cells whose coincidence was above a 95% probability threshold (in practice, this was approximately 15-20% coincidence, which excluded no more than one or two low-firing cells per session).

Across all sessions, we isolated between 137 – 256 PMd and 55 – 93 M1 neurons for Monkey C, and 66 – 121 PMd and 26 – 51 M1 neurons for Monkey M. For the neural covariance analysis, we excluded cells with trial-averaged firing rates of less than 1 Hz. Our GLM models were by necessity poorly fit for neurons with low firing rates. Thus, for the GLM analyses, we only considered neurons with a trial-averaged mean firing rate greater than 5 Hz. Pooled across all monkeys and CF and VR sessions, this gave a population of 918 M1 and 2221 PMd neurons. Given the chronic nature of these recordings, it is certain that some individual neurons appeared in multiple sessions. However, our analyses primarily focus on the population-level relationships which we found to be robust to changes in the exact cells recorded, so we do not expect our results to biased by partial resampling.

### Single neuron covariance analysis

For each trial, we considered all time points between target presentation and successful target acquisition. We binned the neural spiking activity in 10ms bins. We divided each session into five blocks of trials: two from the Baseline, two from Force, and then one from the Washout epochs. In each block, we averaged across trials for each target direction. We then computed the covariance between the spiking activity of all pairs of simultaneously recorded neurons.

Next, we sought to summarize the similarity of this covariance within each block to the Baseline condition to look for learning-related changes. For each pair of neurons, we compared the covariance in the Force and Washout blocks to the Baseline covariance. Across the recorded population on each session, we computed the Pearson’s correlation value to quantify the similarity in the covariance. We also created a reference distribution using correlations within the Baseline epoch to assess the ceiling for this metric when the covariance structure should not have changed. We randomly subsampled the Baseline trials and compared the correlations between each set. We repeated this procedure 100 times for each of the 9 CF and 7 VR sessions. The white distributions in Figure S3 show the correlations for these Baseline trials. For the heat maps shown in Figure S2, we used a simple hierarchical clustering algorithm to sort the neurons in the Baseline condition. To enhance this clustering and visualization, we normalized each row to range from −1 to 1. The same sort order was used in the heat map for each block as a means of visually assessing the consistency in the correlation structure.

### Dimensionality reduction

We counted spikes in 50 ms bins and square root transformed the raw counts to stabilize the variance^14^. We then convolved the spike train of each neuron for each trial with a non-causal Gaussian kernel of width 100 ms to compute a smooth firing rate. We used Principal Component Analysis (PCA) to reduce the smoothed firing rates of the neurons in each session to a small number of components for M1 and PMd separately^14^. PCA finds the dominant covariation patterns in the population and provides a set of orthogonal basis vectors that capture the population variance. We call these covariance patterns “latent activity.” Importantly, this latent activity reflected the firing of nearly all individual neurons.

For the null and potent subspace analysis described below, we needed to select dimensionalities for M1 and PMd. We adapted a method developed by Machens et al^34^ to estimate the dimensionality of our recorded populations. In brief, PCA provides an orthogonal basis set (the principal axes) with the same dimensionality as the neural input. However, the variance captured by many of the higher dimensions (with the smallest eigenvalues) is typically quite small. We estimated the noise in the neural activity patterns using the trial-to-trial variation in the activity of each neuron. We subtracted the activity of each neuron between random pairs of trials for each reach direction. This gave an estimate of the variation in spiking of each neuron (trial-to-trial noise) across targets. We then ran PCA on the neural “noise” provided by this difference for all targets. We repeated this 1000 times, giving a distribution of eigenvalues for each of these noise dimensions. We used the 99% limit of these distributions to estimate the amount of noise variance explained for each dimension. This allowed us to put a threshold on the amount of variance that could be explained by noise. The dimensionality was thus defined by the number of dimensions needed to explain 95% of the remaining variance (Figure 2b shows all sessions for M1 and PMd). Importantly, the dimensionality we estimated was robust to the number of recorded neurons since it reflected population-level patterns. We performed a control where we repeated the above analysis with random subsamples of neurons, taking fixed and equal numbers of neurons from the M1 or PMd populations for a session (Figure S4d) and observed no change in the estimated dimensionality.

### Potent and null subspace calculation

Using the above method, we estimated the dimensionality of the M1 and PMd populations on each session. Since we consistently identified a larger dimensionality for PMd than M1, there existed a “null space” in PMd, which encompassed PMd activity with no net effect on M1^16^. To identify the geometry of the null and potent subspaces, we constructed multi-input multi-output (MIMO) linear models *W* relating the *N* dimensions of PMd latent activity to the *R* dimensions of M1 latent activity (with *N* > *R*):

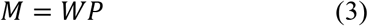

where *M* is an *R* × *t* matrix whose rows contain the *R* dimensions of M1 latent activity, and *P* is an *N* × *t* whose rows contain the *N* dimensions of PMd latent activity; each column of both matrices contains the activity at time *t*. The matrix W, which contains the linear model that maps the PMd latent activity onto the M1 latent activity, has dimensions *R* × *N*.

We then performed a singular value decomposition (SVD) of the matrix *W*:

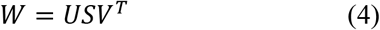

SVD decomposes the rectangular matrix *W* into a set of orthonormal basis vectors that allows us to define the null and potent subspaces. For our purposes, the vectors making up matrix *V^T^* define the potent and null subspaces, with the first *N* rows corresponding to the potent subspace, and the remaining *R* − *N* rows defining the null subspace (Equation 5):

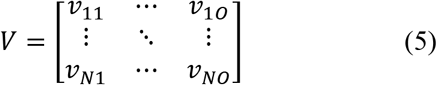

We used only trials from the Baseline period of each session to find the axes for PCA, as well as the null and potent subspaces. The Baseline trials were independent of the CF/VR trials used for both testing and training the GLM models, ensuring that we did not bias our results to find any specific solutions. This is potentially quite important, as it eliminates the possibility that the null and potent spaces simply reflect activity patterns developed during adaptation. In practice, we obtained nearly identical results if we used all of the data, or data only from the CF/VR trials to compute the potent and null subspaces, indicating that they did not change throughout the session. It is also important to note that the null and potent subspaces, as with the PCA axes, typically comprised population-wide activity patterns, rather than sub-groups of neurons. To show this, we defined an index quantifying whether a cell was more strongly weighted towards the output-potent or output-null dimensions. When projected into the potent and null subspaces, the firing rate of each individual cell is multiplied by weights defined in the PCA matrix as well as the matrix defining the null and potent subspaces (*V* in Equation 5). We thus multiplied these two matrices to get an effective weighting of each cell onto the axes of the potent and null subspaces. We then computed an index defined as the difference between the magnitude of the potent and null weights (summed across all dimensions) divided by the total weights. Thus, a value of −1 indicates that the neuron contributed only to null dimensions, +1 indicates the neuron contributed only to potent dimensions, and values near zero indicate the cell has no preference. The resulting distribution, centered on zero, is plotted in Figure S4c.

### Calculating the lag between PMd null and potent activity modulation

We used Canonical Correlation Analysis (CCA) to compare the latent activity in the potent and null subspaces of PMd. In brief, CCA finds linear transformations, that applied to two sets of latent activity, maximize their pairwise correlation^39,66^. Thus, it provides a principled measure of the similarity of signals in the potent and null subspaces. We used CCA to estimate the delay between the null and potent latent activity during Baseline trials within a motor planning window of duration 700 ms following target presentation. We shifted the potent latent activity relative to the null activity by a series of lags (−350 to 350 ms in 10ms steps) and performed CCA at each lag. We identified the lag that maximized the correlation between the activity in each subspace. We repeated this analysis using a block of trials from the end of Force to determine if the lags changed as a consequence of learning (Fig. 6d).

### Generalized Linear Models

In our analyses, we used GLMs to predict the spiking activity of single neurons based on the activity of the remaining population, as well as kinematic signals. We trained Poisson Generalized Linear Models^67^ (GLMs) to predict the spiking activity of individual neurons on a single-trial basis^32^. GLMs extend Gaussian multilinear regression approaches to the Poisson statistics of neural spiking. We took weighted linear combinations of the desired covariates, *_i_*, such as population spiking:

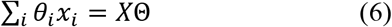

where *X* and *Θ* are matrices containing all *x_i_* and *θ_i_*, respectively. The weighted covariates were passed through an exponential inverse link function. The exponential provides a non-negative conditional intensity function *λ*, analogous to the firing rate of the predicted neuron:

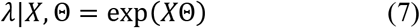

The number of observed spikes, *n*, in any given time bin is assumed to a Poisson process with an instantaneous firing rate mean of *λ*:

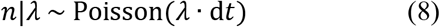

### Covariate inputs to the GLMs

We binned the neural spikes at 50ms intervals and downsampled the continuous kinematic signals to 20 Hz to match the binned spikes and dimensionality-reduced latent activity. We shifted the kinematic signals backwards in time by three bins (150 ms) to account for transmission delays between cortical activity and the motor output. Previous studies have observed a broad range of delays^68^, so we convolved the kinematic signals with raised cosine basis functions centered at 0 ms and −100 ms, adapting the method of Pillow et al., where bases further back in history become wider^31^. By including these convolved signals as inputs to our GLM models, we allowed the neurons to have more flexible temporal relationships with the kinematics. Note that all GLM models in the main text included the same convolved endpoint position, velocity, and acceleration signals as covariates. For Figure S7c,d, we performed a control in which we repeated the GLM models for the CF task using only the velocity inputs.

We trained two types of models: the Basic models included only kinematic covariates, while the Full models included both the kinematic covariates and the spiking activity of the neural populations (Figure S5a). For the GLM analysis with neural population latent activity as inputs, we selected a dimensionality for M1 and chose the PMd dimensionality to be twice this value. This decision was to control for the number of inputs to the GLM when analyzing the null and potent space activity. For the **PMd**-M1 model, we identified the low-dimensional latent activity in PMd (see above) and used these as inputs to the GLM. For **M1**-M1 and **PMd**-PMd, we left out the single predicted neuron and computed the latent activity using the remaining neurons in that brain area. We then used these signals to predict the activity of the left-out neuron. For **Pot**-M1 and **Null**-M1, we projected the time-varying PMd activity onto the axes for the potent and null subspace, respectively (see above). We then used these time-varying signals as inputs to GLMs to predict the spiking of M1 neurons. For the GLMs with single neuron inputs, we trained three different types of Full models. **M1sn**-M1 models predicted the spiking activity of each M1 neuron from the activity of all other M1 neurons recorded on the same session, **PMdsn**-PMd models predicted the spiking of each PMd neuron from all other PMd neurons, and **PMdsn**-M1 models predicted M1 neurons using the activity of all PMd cells. Since PCA captures population-wide covariance patterns, we expected that this approach would provide nearly identical results to the single neuron models, and it was included primarily as a control.

### Training the GLMs

We trained the models using the last 50% of Force or Rotation when behavior was most stable, including only trials in which the monkeys reached successfully to the outer target (reward trials) (Figure S5d). This allowed us to test the generalization of the GLMs between late and early adaptation trials. For the CF, it was important to both train and test the GLMs using trials from the Force epoch to avoid extrapolating between the Null and CF conditions: when we imposed the CF, it changed the relationship between the kinematics and dynamics of limb movement. Thus, if we trained the GLM on Baseline trials, the altered relationship between kinematics and neural activity during CF trials^28^ would lead to poor GLM generalization for all models. By both training and testing within the Force epoch, we avoided the problem of extrapolating to new dynamics conditions. Although the VR sessions did not have this problem, we adopted this same approach for the sake of consistency. We performed an additional analysis to look for deadaptation effects in Washout (Figures 5,7f). The procedures were similar for this analysis, though we trained the GLMs using Baseline data and then tested using trials from early Washout.

We trained the models using a maximum likelihood method (glmfit in Matlab, The Mathworks Inc). In the case of our full population spiking models, we had dozens to hundreds of covariate inputs for a single predicted output. Although we had very large numbers of training datapoints (typically on the order of 10,000 samples), there is the possibility our models were impaired by overfitting. We guarded against overfitting using ten-fold cross-validation of our training dataset. We also repeated our analyses using Lasso GLM for regularization and observed nearly identical results (data not shown). We thus chose to use the non-regularized GLM for simplicity and to reduce the computational load, since it did not impact our results.

### Evaluating GLM performance

We evaluated GLM performance using a particular formulation of the pseudo-R^2^ (pR^2^). The pR^2^ is analogous to the R^2^ commonly used in model-fitting with Gaussian statistics, but it is generalized to incorporate the approximate Poisson statistics of the neural spiking data:

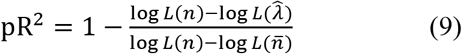

The pR^2^ finds the difference in log-likelihood between the observed spiking data (*n*) and the model predictions 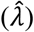. This value is compared against the difference in log-likelihood for the mean of the dataset 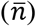. We used the Likelihood (*L*) for Poisson data according to:

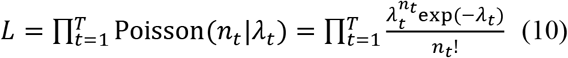

And thus, the log-likelihood (log *L*) across all time bins (t) of a given spike train is:

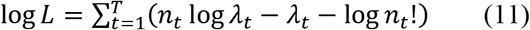

Although the upper bound for pR^2^ is one, poor model fits can be less than zero. A pR^2^ of one indicates a perfect model fit, a value of zero indicates that the model prediction performs as well as finding the mean of the data, while values less than zero indicate that the model performed worse than merely fitting the mean. pR^2^ values are smaller in magnitude than those typically found with the Gaussian R^2^. When evaluating GLM fits on a block of data, we used a bootstrapping procedure to resample the data with 1,000 iterations to obtain 95% confidence bounds on the pR^2^ value. We considered a model fit to be significant if this bootstrapped confidence interval was above zero, indicating that the model helped to explain the spiking activity.

For many analyses, we used the relative pseudo-R^2^ (rpR^2^), which directly compares two separate GLM models. While pR^2^ compared the log-likelihood of the model predictions to the mean of the data, the rpR^2^ compares the predictions of a Full model to a Basic model with fewer covariates^33^.

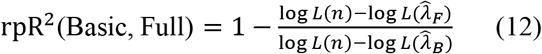

Here, 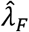, the Full model prediction, which includes both the kinematics and the population spiking, is compared to 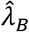, the prediction of the Basic model, which includes only kinematics. This metric thus quantifies the improvement in performance afforded by the additional neuronal inputs. Positive values indicate that the Full model performed better than the Basic model, while negative values indicate that predictions were better with kinematics alone. As with the pR^2^, we obtained confidence bounds with a bootstrapping procedure and assessed significance by determining if the lower bound was above zero. This indicated that the addition of population spiking added information over the kinematics alone, and thus could be capturing statistical dependencies between the population and the predicted cell.

For the time course of GLM model performance plots, such as Figure 3f, we predicted neural spiking on individual trials. However, single-trial predictions could be quite noisy, particularly for neurons with relatively low firing rates. For example, if a cell fired very few spikes on a particular trial, the pR^2^ may be quite low, even though the model otherwise performed quite well when viewed across all trials. To remove some of this variability for purposes of visualization, we smoothed the trial-to-trial predictions for each neuron (together with the overlaid behavior) with a moving average. We chose a window of 30 trials, though we observed similar (but noisier) traces even down to window sizes of 5-10 trials. Since there were rapid behavioral improvements in the early trials, we padded the beginning of the block of trials with NaNs, each of a length of half of the window size. This helped to prevent averaging out the changing behavioral effects, with the tradeoff of increased noise. In practice, the appearance of the figures was similar without this padding.

### Selecting cells with significant population relationships

For most of our analyses, we studied cells that were well-predicted by our GLMs. We determined this by two main criteria using ten-fold cross-validation on the training data. First, we required that the Basic pR^2^ was significantly above zero. This reduced the pool of candidate cells to 522 of 918 (57%) in M1 and 612 of 2221 (28%) in PMd, but was necessary so that the rpR^2^ would be well defined. Qualitatively, we obtained similar results when we relaxed this criterion to include more cells. We also required that the rpR^2^ was significantly above zero. We only included cells that were significantly above zero for all ten of the folds for both pR^2^ and rpR^2^. This conservative method ensured that we only studied cells that were reliably predicted.

### Calculating GLM error

To compare performance of the different GLM models, we first defined a fixed reference point of eight consecutive trials in Late adaptation, which was close to (but independent from) the trials from the second half of adaptation that we used to train the GLMs (around 70 trials from the start). On each trial, we computed the difference in rpR^2^ relative to this Late adaptation performance. To account for inherent differences in performance of the models, we normalized this difference by the rpR^2^ computed from the cross validated training set. Thus, the model error metric represents the % change in performance from Late to Early adaptation. Note that as constructed, the model error is necessarily zero in Late adaptation (such as when plotted in Figure 3g). This metric provides a compact way to compare whether the models performed worse during Early adaptation compared to Late adaptation. For the Washout predictions, the rate of deadaptation was faster than adaptation in either Force or Rotation. Thus, the blocks of trials that we selected for the Early and Late reference when testing the GLMs in Washout were closer to each other in time than the blocks we used for the Force or Rotation epochs.

### Controls for the potent and null subspaces

For the PMd potent and null subspaces, we ensured our results did not depend on the particular dimensionality chosen. We repeated the full GLM analysis selecting different M1 dimensionalities ranging from 5 to 20. We enforced that PMd always had twice the number of dimensions of M1 to control for the number of inputs to the GLM models. We then trained the GLM models using the methods described above and computed the model error for each dimensionality (shown in Figure S7e).

### Statistical tests

For the GLM models, we assessed the significance of model fits empirically using a bootstrapping procedure on cross-validated data as described above. Additionally, we used two-sample statistical tests to compare the distributions of pR^2^ changes in Early and Late adaptation. For Figures 3e and S5f, we used a Kolmogorov-Smirnov test of the rpR^2^ values. For the Model Error comparisons, which summarized differences in the metric, we used a *t*-test of the normalized change in rpR^2^. All tests were two-sided.

## Supplementary Figures

**Figure S1.**
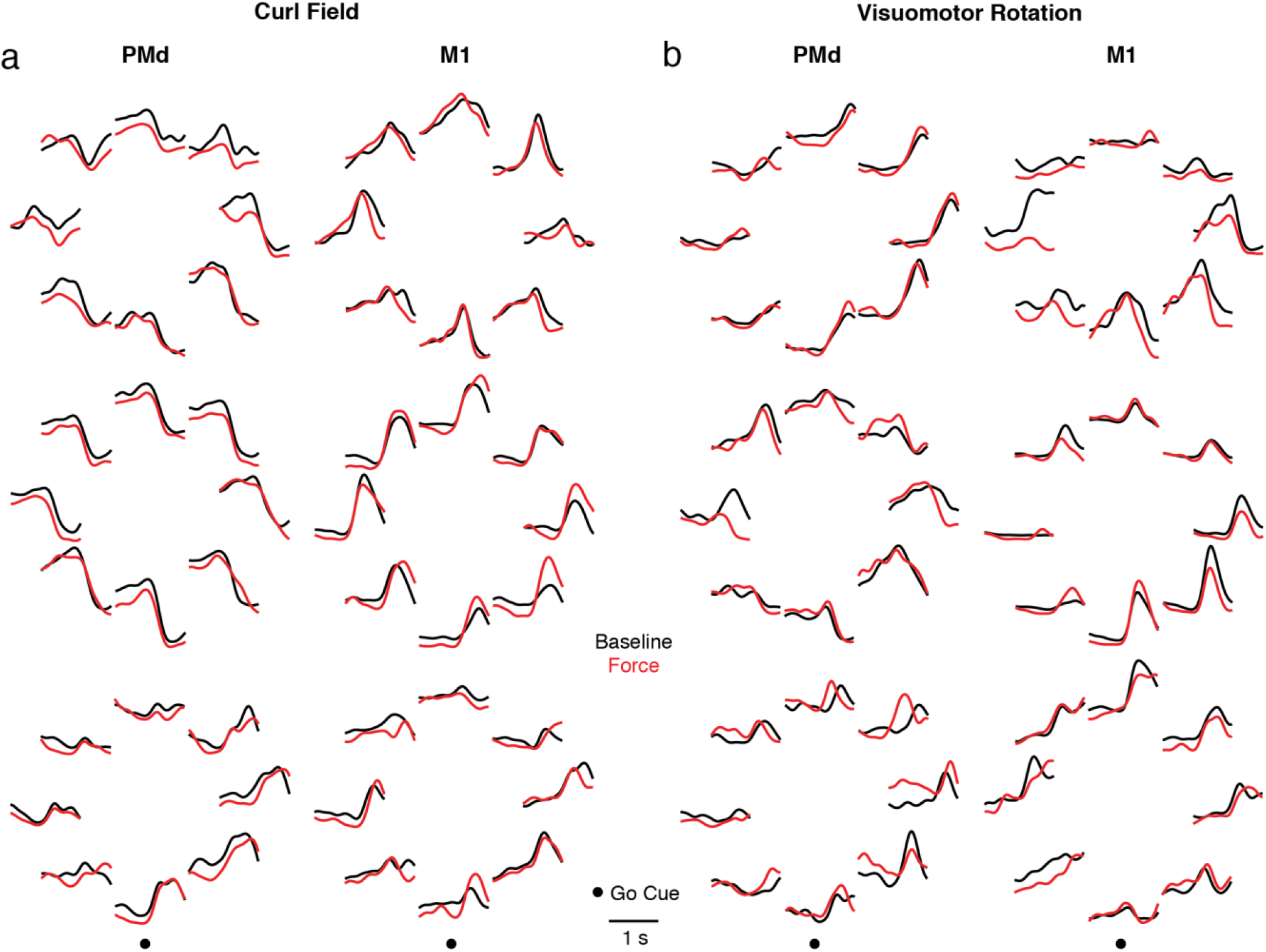
Example neural activity before and after CF and VR adaptation. a) Trial-averaged firing rates for three example PMd (left) and M1 (right) neurons from the same CF session, for each of the eight targets. Neural activity was smoothed with a Gaussian kernel, aligned to the go cue (indicated by the dot at the bottom of each panel), and averaged across reaches to each target during Baseline (black) and end of Force (red). b) Same as Panel a, for a VR session.

**Figure S2.**
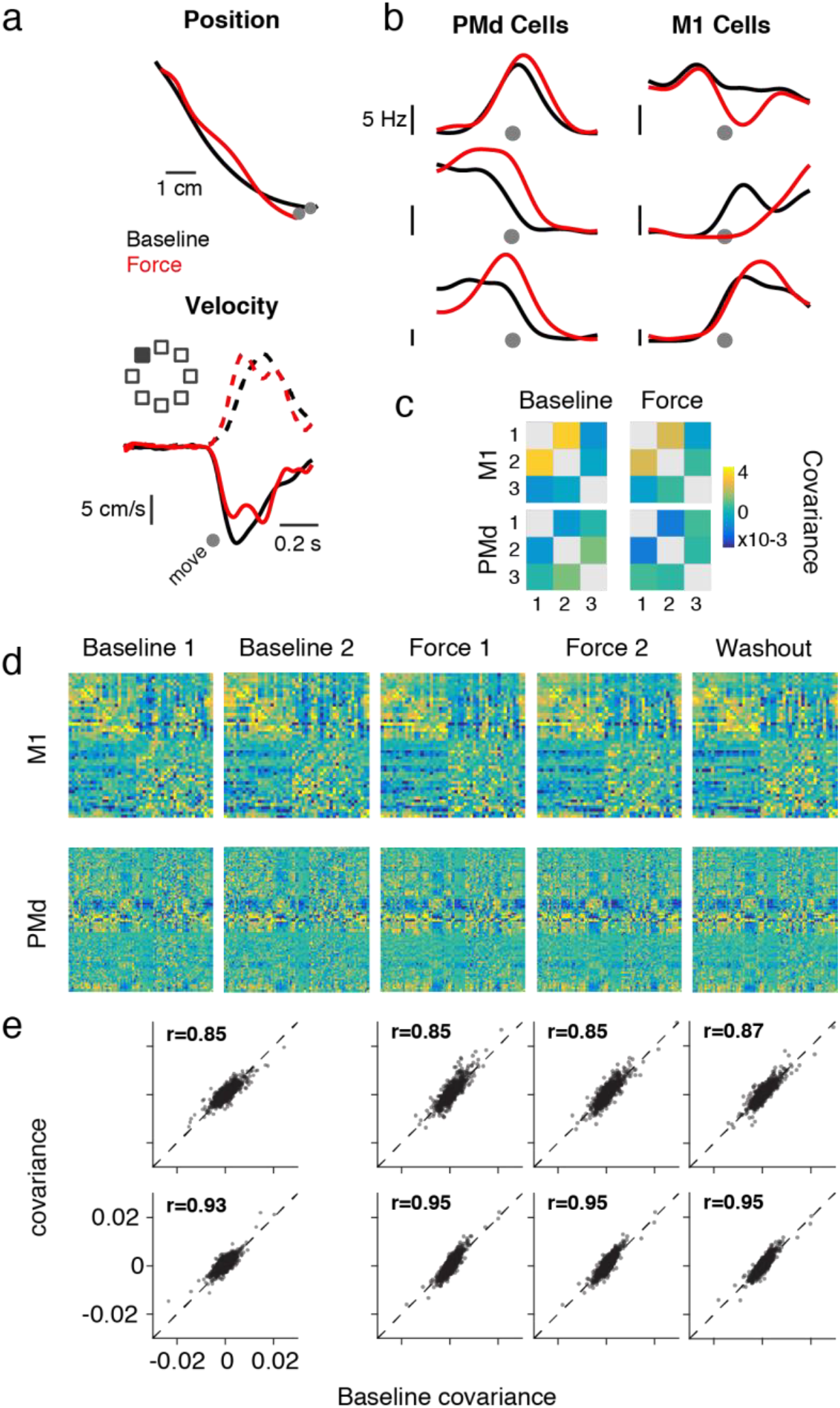
Neural covariance during CF adaptation. a) Example position and velocity traces for reaches in a single target direction in Baseline (black) and Force (red) for horizontal (solid) and vertical (dashed) axes. Traces are averaged across the last three successful reaches to the 135° target in each epoch. b) For the same reaches as Panel a, example firing rates of two M1 neurons and three PMd neurons, where each axis is the firing rate of a neuron and the firing rates are plotted as a function of time next to each axis. c) Covariance matrix relating the example neurons in Panel b before and after adaptation. The covariance was computed using all eight target directions, but only a single direction is plotted in Panels a and b. d) Covariance (see Methods) between all cells recorded on the same session as Panels a-d. We normalized the covariances such that the min and max for each neuron spanned the full dynamic range, clustered the Baseline covariance, and kept the same order for all remaining plots. We compared two halves of Baseline trials, the Washout trials, and split the Force epoch into two blocks to compare to the full set of Baseline trials. The M1 and PMd populations were analyzed and plotted separately. e) Summary of covariance structure between BL and all other blocks for all combinations of neurons recorded in each of the nine CF sessions. The r value for each plot indicates the Pearson’s correlation coefficient between the covariance values for the two task epochs. The covariance was highly consistent throughout the task.

**Figure S3.**
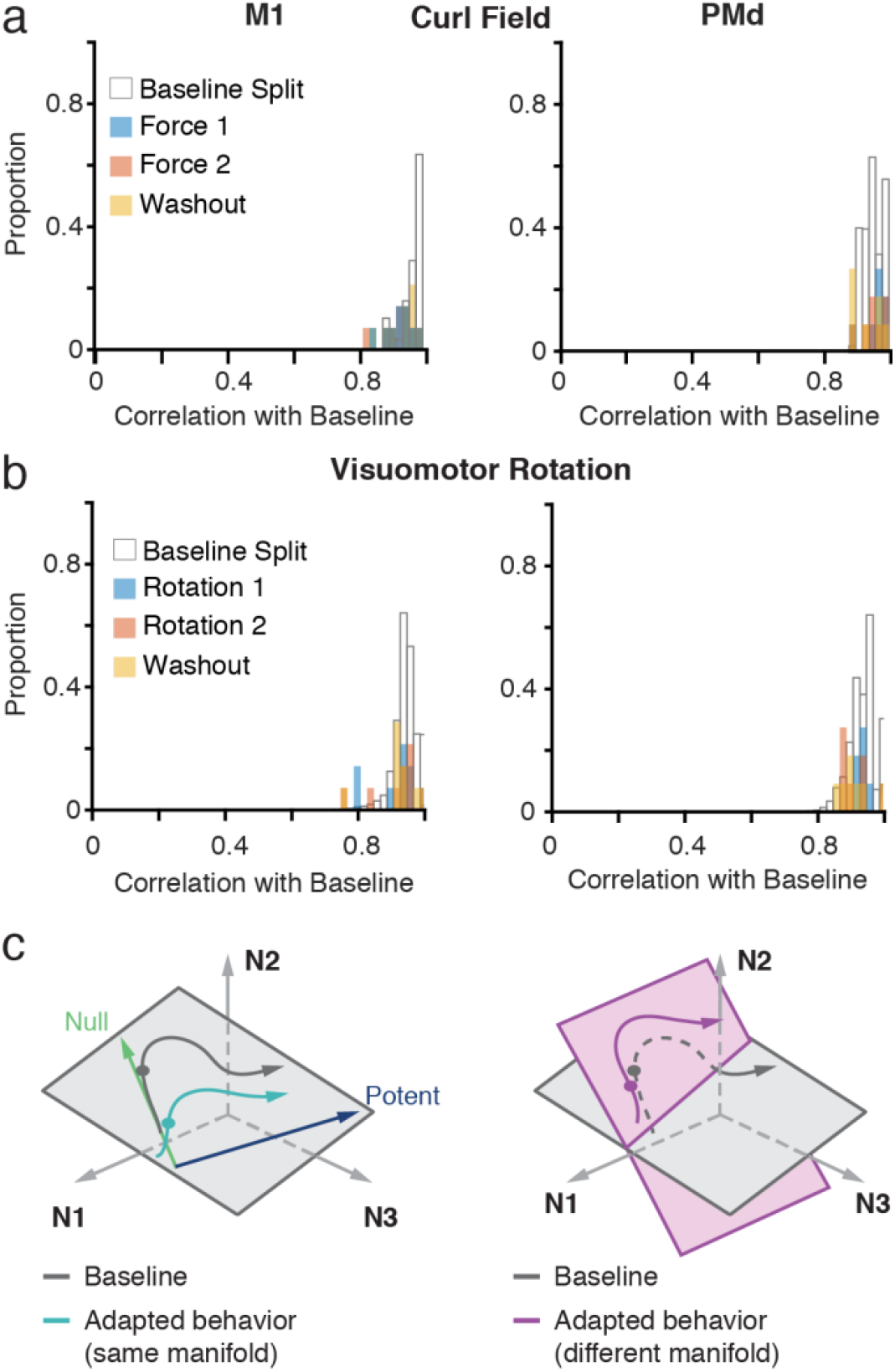
Neural covariance structure during adaptation for all sessions. a) Summary histograms of neural covariance similarity across all CF sessions. Each count in the colored distributions corresponds to the population-wide Pearson’s *r* value, as shown in Figure S2e comparing each of the four blocks of trials to Baseline, as well as comparisons within Baseline (white bars). The real data distributions strongly resemble the split Baseline ones, indicating that the changes in covariance between Baseline and each of the Force and Washout bins are similar to the changes in covariance in Baseline bins. Therefore, there were likely no adaptation-related changes in the population covariance structure. b) Same as Panel a to summarize across all VR sessions. c) In order to correctly interpret the results of our study, it is important to discuss what “changes” occur in the output-null subspace of PMd during CF adaptation. The degradation of the **Null**-M1 GLM model is not likely to be primarily due to a change in the orientation, or “structure”, of the null subspace (see Figure S2, S3 and GLM results). Instead, PMd activity explores this subspace in a nove1 manner. In this cartoon, we highlight the seemingly subtle but crucial distinction between these ideas. On the left panel, we show a 3-D “state space” for three neurons (N1, N2, N3). For a Baseline reach to a single target, the activity of these neurons follows the black line as it evolves from movement preparation along the null axis, and finally to movement initiation along the potent axis. This activity is confined to a 2-D manifold (gray plane). After CF adaptation, the neural activity takes a different path (blue). However, critically, activity both before and after learning lie within the same Baseline manifold, which has the same potent and null subspaces. The right panel offers an alternative possibility. The same Baseline reach and manifold is shown in gray. However, during learning the orientation of the 2-D manifold could change (purple plane). Neural activity in the adapted reach is now constrained to this new manifold. Our results are best viewed with the first (left panel) hypothesis: our within-area GLM models predicted well, and the covariance structure in M1 and PMd did not change. These observations suggest that the neural covariance defining these manifolds was also unchanged. Thus, as our monkeys learned, the null activity in PMd explored a fixed null subspace.

**Figure S4.**
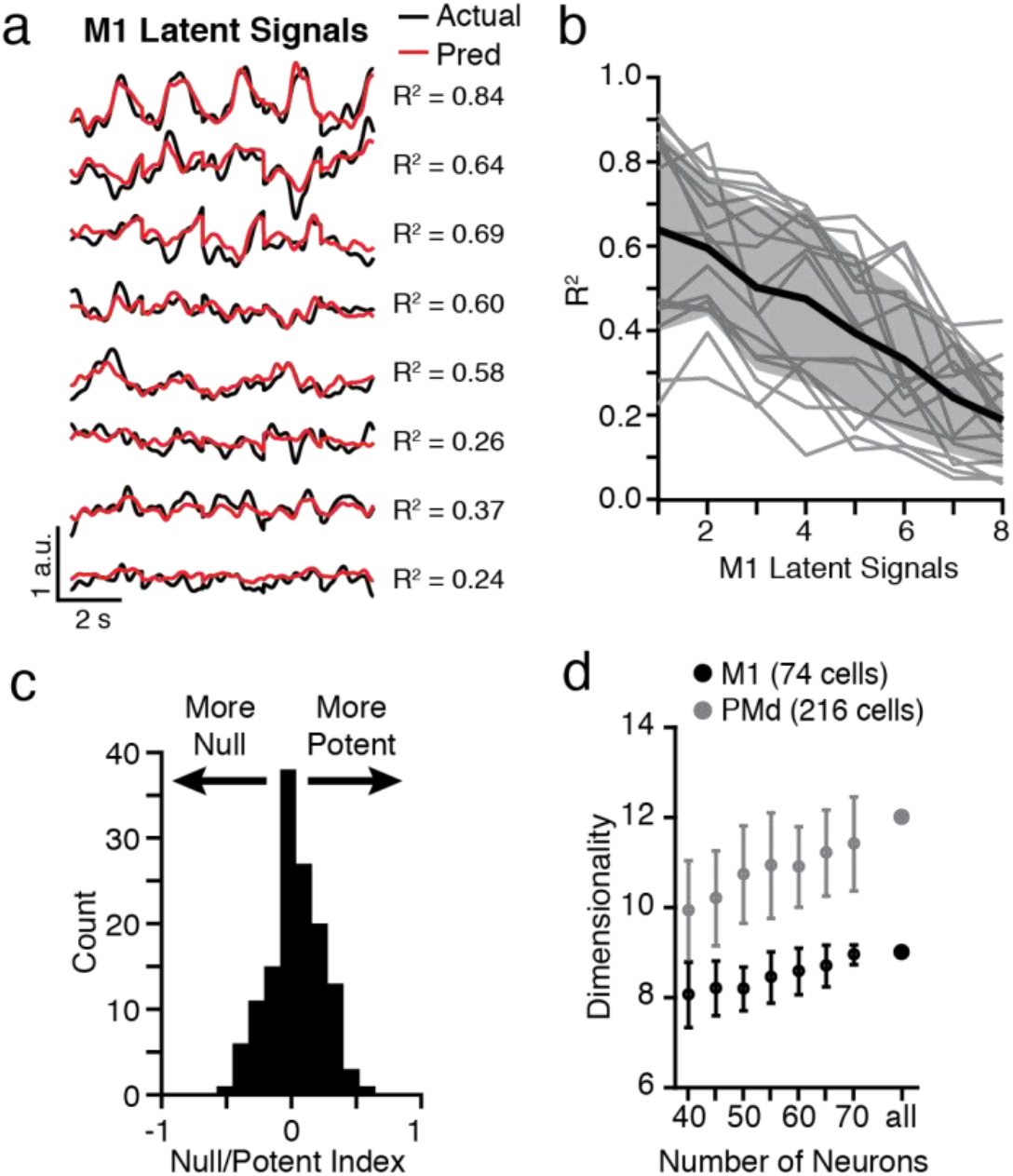
Calculation of the PMd null and potent subspaces. a) Example predictions (red) of the first eight principal components of M1 latent activity (black) from the first sixteen principal components of PMd latent activity, with R^2^ quantifying quality of fit for a single session. b) Summary of R^2^ for predictions of each of the eight principal components of M1 latent activity across sessions (gray lines). Black line and shaded area indicate and mean and standard deviation across sessions. c) We attempted to identify potent or null subpopulations of neurons using an index that quantified the relative weights of each neuron onto the potent and null axes (see Methods). Values of 1 indicate the cell was exclusively potent, and values of −1 indicate the cell was exclusively null. The distribution of cells was centered around zero and virtually no cells had a null/potent index close to −1 / 1, indicating that the potent and null subspaces captured population-wide features. d) The effect of population size on dimensionality for one example session from Monkey C. We randomly subsampled the neural populations 100 times at fixed neuron counts and repeated the dimensionality analysis. The result of each repetition is plotted as a mean and standard deviation. PMd (gray) was consistently higher dimensional than M1 (black).

**Figure S5.**
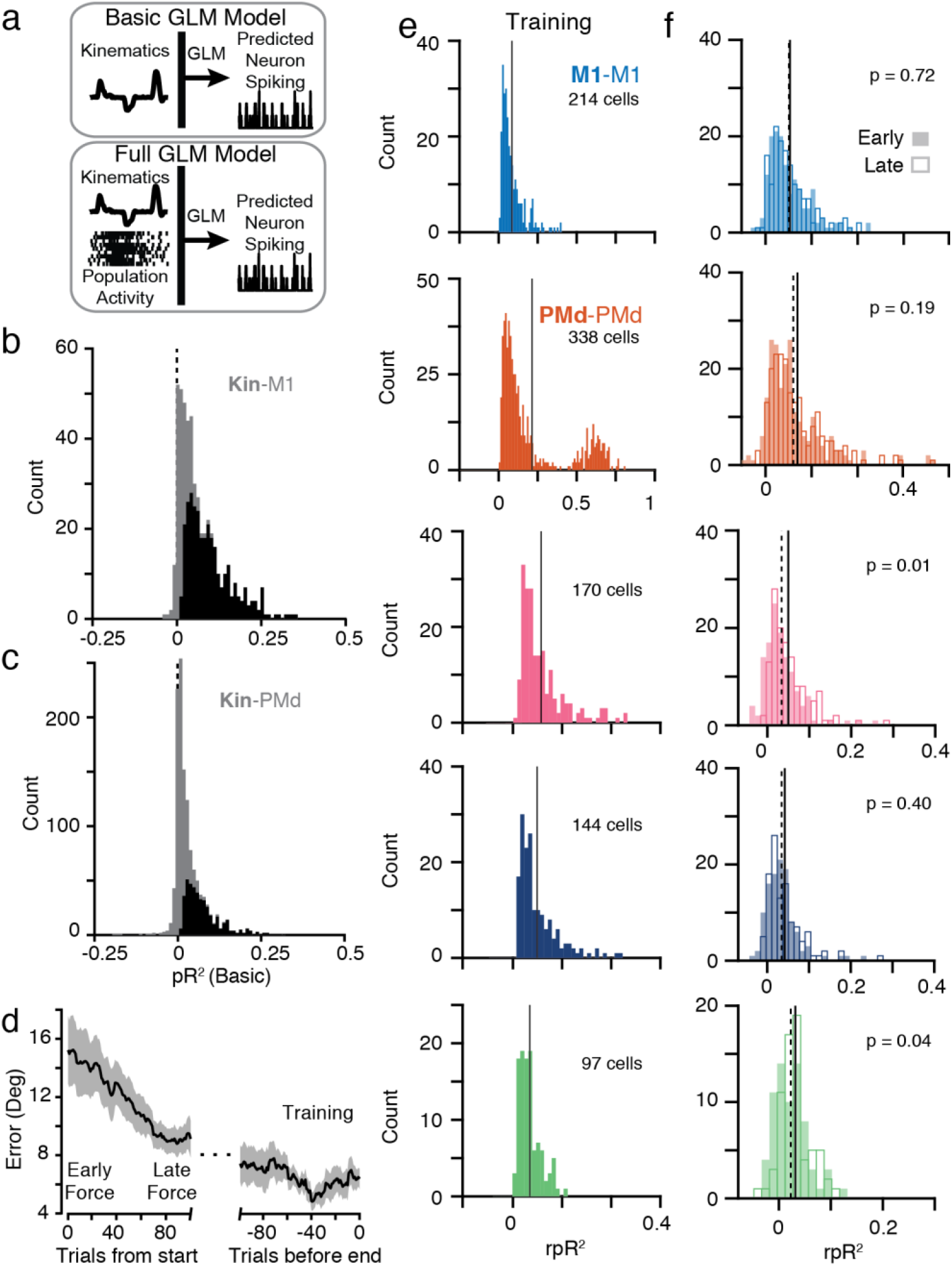
GLM model performance. a) Schematic representation of the GLM models. The Basic model included only kinematic covariates (see Methods), while the Full model included both kinematics and neural activity. The relative pseudo-R^2^ metric was a comparison between these two models. b) Distribution of cross-validated pseudo-R^2^ values for predictions of all M1 neurons from all sessions with the Basic model (gray). Black overlaid distribution shows cells with significant model fits selected for consideration in the later analyses (see Methods). c) Same as Panel b, but for predictions of PMd neurons. d) Illustration of the training and testing process. Lines indicate the mean and standard deviation of behavioral error during CF learning aligned on the first CF trial (left) or the last CF trial (right). The GLMs were trained using the end of learning data and tested during the earlier adaptation period. e) Distribution of cross-validated relative pseudo-R^2^ values for the five GLM models. f) Distribution of unprocessed rpR^2^ values for all predicted cells for the five GLM models during Early and Late CF adaptation. Only **Null**-M1 and **PMd**-M1 were significantly changed between the two periods.

**Figure S6.**
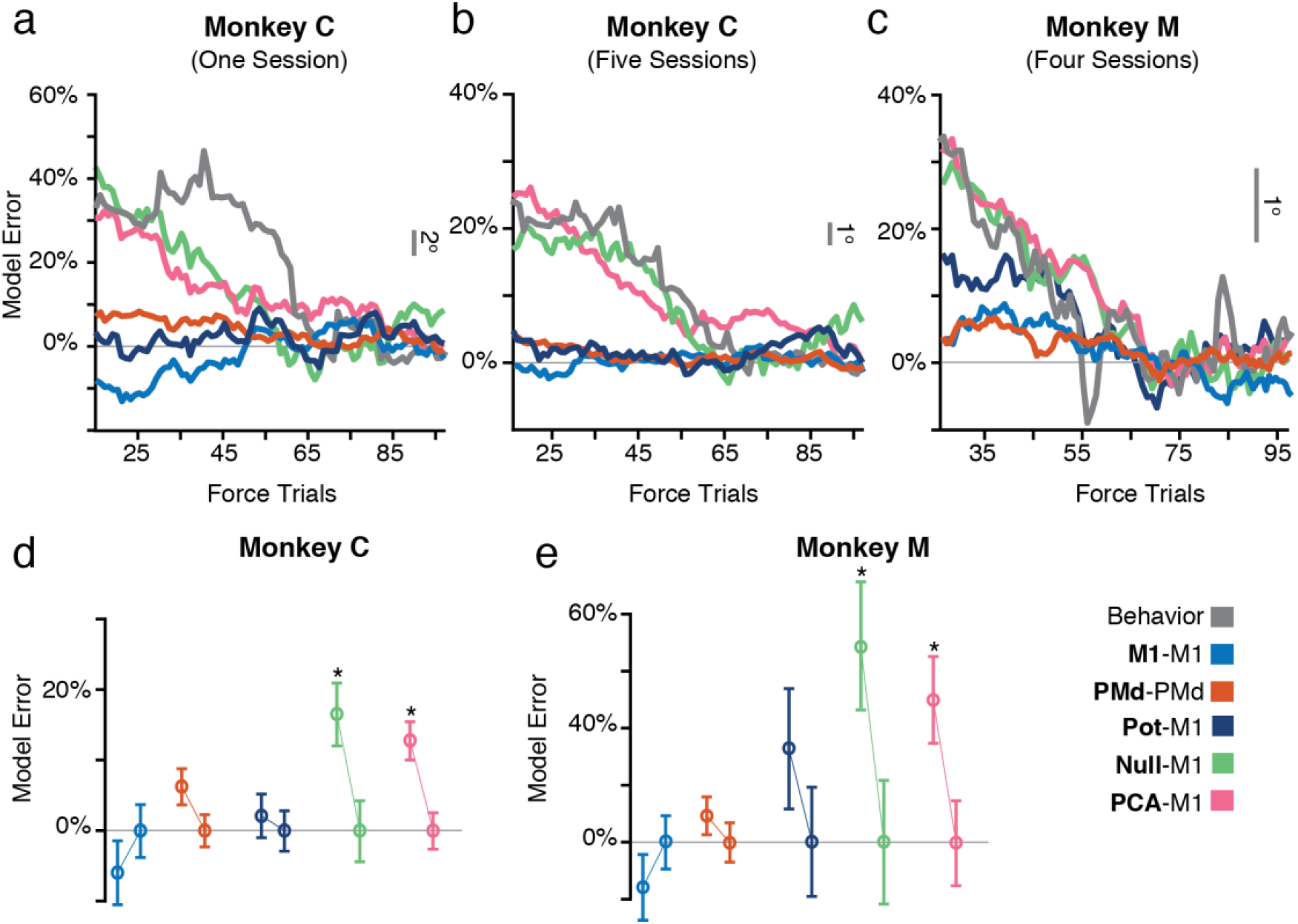
Monkey-specific GLM model performance. a) Formatted as Figure 3f, but for all neurons recorded on a single session from Monkey C. The gray line shows a moving average behavioral error of the monkey on that session for the same trials. Note that this is the same session from Monkey C as the previous example data, such as Figure S2. b-c) Same as Panel a, but for all sessions from Monkey C (b) and Monkey M (c). Since Monkey M contained fewer neurons, predictions were considerably noisier and we had fewer predicted neurons to average over. Thus, we extended the moving average window size to 50 trials to better illustrate the time course. d) Summary of model error distributions for Monkey C alone. Plotted are mean and standard error for Early (left) and Late (right) trials for each of the five GLM models used. Asterisks indicate significant differences between the Early (left connected bar) and Late (right connected bar) trials at p < 0.01 (two-sample t-test). e) Same as Panel d, but for Monkey M alone.

**Figure S7.**
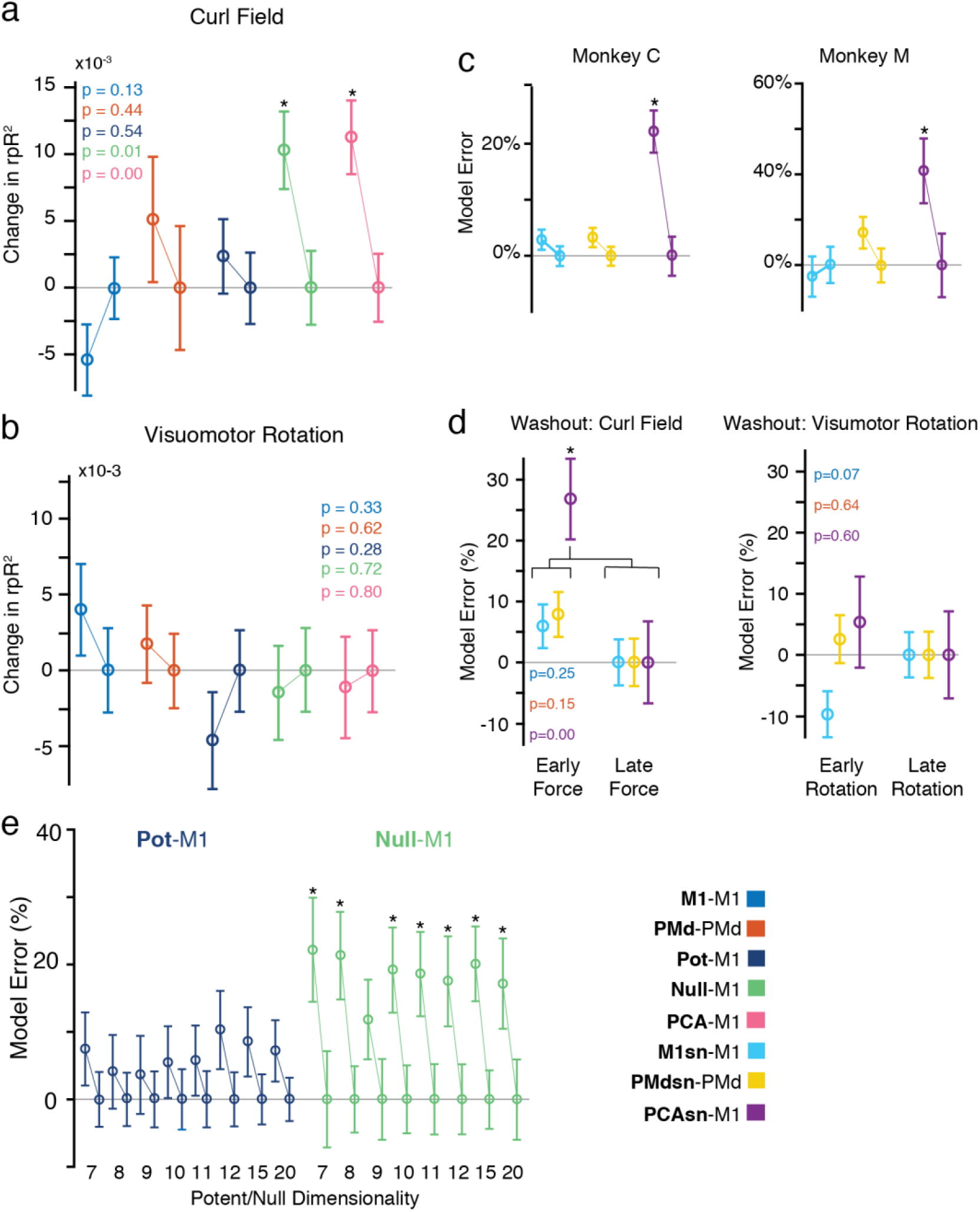
GLM controls. a) The distributions of change in rpR^2^ for the curl field sessions, but without the normalization by the cross-validated performance. The same significant effects are observed. b) Same as Panel a, but for the visuomotor rotation sessions. There were no significant differences in the models. c) We repeated the GLM analysis using single neuron spiking as inputs, rather than the dimensionality-reduced population activity. We computed models for M1 predicting M1 (**M1**-M1), PMd predicting PMd (**PMdsn**-PMd), and PMd predicting M1 (**PMdsn**-M1). The Model Error distributions are summarized here for Monkey C (left) and Monkey M (right). Consistent with the results of Figures 3 and 4, only **PMdsn**-M1 generalized poorly. d) Testing the Washout epoch performance for the single neuron models. The effect of de-adaptation is observed in the curl field task for **PMdsn**-M1 only, and no change was observed in the visuomotor rotation task. e) Comparison of GLM performance error between early CF (left bars) and late CF trials (right bars) for **Pot**-M1 and **Null**-M1 as a function of the selected dimensionality. Asterisks indicate significance at p < 0.01 (two-sample t-test) for comparison between Early (left) and Late (right) for each model. The results for a dimensionality of eight plotted here are the same as in Figure 3d. Our primary effect that **Pot**-M1, and not **Null**-M1, generalizes to early CF trials held for a wide range of dimensionalities.

